# Supercharging carbohydrate-binding module alone enhances endocellulase thermostability, binding, and activity on cellulosic biomass

**DOI:** 10.1101/2023.09.09.557007

**Authors:** Antonio DeChellis, Bhargava Nemmaru, Deanne Sammond, Jenna Douglass, Nivedita Patil, Olivia Reste, Shishir P. S. Chundawat

## Abstract

Lignocellulosic biomass recalcitrance to enzymatic degradation necessitates high enzyme loadings incurring large processing costs for industrial-scale biofuels or biochemicals production. Manipulating surface charge interactions to minimize non-productive interactions between cellulolytic enzymes and plant cell wall components (e.g., lignin or cellulose) via protein supercharging has been hypothesized to improve biomass biodegradability, but with limited demonstrated success to date. Here we characterize the effect of introducing non-natural enzyme surface mutations and net charge on cellulosic biomass hydrolysis activity by designing a library of supercharged family-5 endoglucanase Cel5A and its native family-2a carbohydrate binding module (CBM) originally belonging to an industrially relevant thermophilic microbe *Thermobifida fusca*. A combinatorial library of 33 mutant constructs containing different CBM and Cel5A designs spanning a net charge range of −52 to 37 was computationally designed using Rosetta macromolecular modelling software. Activity for all mutants was rapidly characterized as soluble cell lysates and promising mutants (containing mutations either on the CBM, Cel5A catalytic domain, or both CBM and Cel5A domains) were then purified and systematically characterized. Surprisingly, often endocellulases with mutations on the CBM domain alone resulted in improved activity on cellulosic biomass, with three top-performing supercharged CBM mutants exhibiting between 2–5-fold increase in activity, compared to native enzyme, on both pretreated biomass enriched in lignin (i.e., corn stover) and isolated crystalline/amorphous cellulose. Furthermore, we were able to clearly demonstrate that endocellulase net charge can be selectively fine-tuned using protein supercharging protocol for targeting distinct substrates and maximizing biocatalytic activity. Additionally, several supercharged CBM containing endocellulases exhibited a 5–10 °C increase in optimal hydrolysis temperature, compared to native enzyme, which enabled further increase in hydrolytic yield at higher operational reaction temperatures. This study demonstrates the first successful implementation of enzyme supercharging of cellulolytic enzymes to increase hydrolytic activity towards complex lignocellulosic biomass derived substrates.

## INTRODUCTION

As demand for fossil-fuel based petroleum products and fuels continues to increase, global oil production is forecasted to reach its peak^1^. Resource scarcity and climate change exasperated by this petroleum dependance requires suitable sustainable energy sources in order to replace conventionally derived fuels and chemicals and promote a circular bioeconomy^2–4^. Bioethanol is one potential candidate to replace conventional fuels that can be produced from abundant carbon neutral renewable sources like biomass^5,6^. In order to become an economically feasible alternative energy that can compete with the price of fossil fuels, biorefineries must valorize lignocellulosic wastes into useful fuels and chemicals^7,8^. Lignocellulosic waste biomass is a readily abundant and underutilized resource^9^ that can be derived from agricultural residues like corn stover or sugar cane bagasse, as well from woody forest products^10^. These substrates are rich in insoluble polysaccharides like cellulose and hemicellulose that form a tightly packed hydrogen bonded network within the secondary cell walls of plant masses buried within a layer of the structural polymer lignin^11^. These complex polysaccharides are subject to enzymatic saccharification to their fermentable monomers for biofuel production by Carbohydrate Active enZymes (CAZymes) that catalyze the hydrolysis of glycosidic linkages within glucan chains. There are numerous different CAZyme families with mechanistic differences and substrate specificities that can act synergistically to completely depolymerize the biomass complex^12–15^. However, this is an idealized scenario, and real-world biomass conversion to biofuels is inefficient due to limited technologies and processing challenges related to the substrate.

The economic viability of biofuel production from lignocellulosic biomass is significantly hindered by biomass’ recalcitrance to enzymatic saccharification which is a large contributor to high processing costs^16^. The tight biomass structure within plant secondary cell walls provides limited access for enzymes to depolymerize cellulose and hemicellulose significantly reducing the ability to hydrolyze these polysaccharides^17^. Cellulose itself exists as insoluble crystalline microfibrils which poses a challenge for enzymes to catalyze hydrolysis at the solid-liquid interface^16^. Lignin is also known to contribute to biomass recalcitrance^16,18^ through the irreversible non-productive binding CAZymes^19^. These factors greatly reduce the efficiency of enzymatic saccharification resulting in high enzyme loadings to supplement low activity and enzymes losses, which greatly increases the processing costs in biorefinieries^20^. One solution to this issue is the introduction of biomass pretreatment prior to enzymatic saccharification in order to make cellulose and hemicellulose more accessible to CAZymes reducing overall recalcitrance^21,22^. Thermochemical pretreatment methods like steam explosion, ammonia fiber expansion (AFEX), and extractive ammonia (EA) are successful in disrupting the biomass matrix, and are even capable of converting the native cellulose-I into a more digestible allomorph significantly reducing recalcitrance^23,24^. However, non-productive binding remains a persistent issue as these methods do not totally remove lignin, and pretreatment does not do much to supplement low activity on crystalline cellulose^16,25^. Many of the pertinent CAZymes exist as multifunctional^15^ and multimodular proteins containing a carbohydrate binding module (CBM) and catalytic domain (CD) linked through a flexible linker peptide^26,27^. One approach to overcome the limitations that persist from pretreatment alone is to utilize the rational design and engineering^28^ of CBMs to modulate productive vs. non-productive binding interactions, as well as engineer CDs with higher catalytic activity.

Evidence suggests that there is significant surface charged interactions between lignin, cellulose, and CAZymes. Binding studies of different green fluorescent protein (GFP) mutants binding to lignin confirm a weak correlation between increasing positive net charge and greater irreversible binding to lignin. This relationship likely stems from electrostatic interactions between the slightly negative lignin surface and charged proteins^29^. Applying this principle to a family-5 glycosyl hydrolase CelE CD appended to a family-3a CBM, Whitehead et al. (2017) were able to create lignin resistant cellulases^30^. Utilizing protein supercharging^31^ to introduce aspartate and glutamate mutations in solvent exposed amino acid residues, the introduced negative surface charges successful reduced lignin inhibition at the expense of overall catalytic activity on amorphous phosphoric acid swollen cellulose (PASC)^30^. This result provides insight that surface charged interactions can indeed reduce binding to lignin but can also impact binding and subsequent hydrolysis of cellulose model substrates that carry a similar negative surface charge. This work provides promise in using protein surface supercharging for improving cellulose hydrolysis, but several questions still remain unknown. These include (i) what effect does positive supercharging have on both CBM and CD, (ii) how do supercharged cellulases behave on crystalline cellulose and biomass, and (iii) is there a specific net charge where catalytic turnover is maximized on different substrates. The last question also aligns with the Sabatier principle^32^ that has been previously applied to cellulases^33,34^ and suggests that intermediate binding interactions between substrate and enzyme provide optimal catalytic turnover. Applying this principle to supercharged enzymes, it may be possible to increase/decrease binding between enzyme and substrate by tweaking surface charged interactions so that at a critical net charge one would observe optimal catalytic turnover.

Here we build upon the knowledge of protein supercharging’s effect on cellulose hydrolysis by supercharging a family-5 endocellulases Cel5A and its native family 2a CBM from the thermophilic microbe *Thermobifida fusca*^35,36^. Rosetta macromolecular software was used to supercharge both CBM2a and Cel5A CD for a total library of 33 mutants (including wild type full length enzyme) spanning a net charge range of −52 to 37. Mutant activity was screened first from soluble cell lysates, with those constructs that performed best targeted for large scale expression and purification. Purified enzyme assays identify a much greater contribution to activity improvements when the binding module alone was engineered compared to the CD. This result is likely linked to improved binding affinity for supercharged CBMs elucidated by GFP based pull down binding assays. Furthermore, hydrolysis assays screened at different solution pH identify some charge optima that exist on biomass and cellulose, as well as shifts in optimal pH for supercharged mutants. Through studying hydrolytic activity at elevated temperatures, positively supercharged mutants were found to exhibit an increased optimal hydrolysis temperature and resulting increase in hydrolytic yield on crystalline cellulose substrates. There are several mutants within this supercharged Cel5A library capable of improved catalytic activity compared to wild type enzymes with upwards of 2-fold improvements in activity.

## EXPERIMENTAL SECTION

### Reagents

AFEX pretreated corn stover used for hydrolysis assays were prepared and provided by Dr. Rebecca Ong’s lab (Michigan Technological University) following established protocols^23^. Crystalline cellulose-I was procured from Avicel (PH-101, Sigma-Aldrich), and was also used to prepare phosphoric acid swollen cellulose (PASC) from prior protocols^37^. Chromogenic para-nitrophenyl cellobioside (pNPC) was obtained from Biosynth. All genes used for expression of recombinant constructs were provided by the Department of Energy Joint Genome Institute (DOE-JGI). All other reagents used were purchased from either Sigma Aldrich or Fischer Scientific unless otherwise noted in subsequent sections.

### Computational design of mutant enzyme libraries and plasmid construction

Protein supercharging was done by introducing either positive (K and R) or negative (D and E) amino acids on the surface of either CBM or CD using Rosetta macromolecular software. The structure of CBM2a from *T. fusca* has yet to be solved, thus CBM designs were based off of homology model constructed using Rosetta CM^38^. Supercharged designs for the Cel5A catalytic domain were constructed based off of solved crystal structure (PDB: 2CKR). In a previous study from our group, FoldIt interface was used to identify folding energy change caused due to mutations of individual residues^30^. However, this protocol is not amenable to automation when large mutant libraries are being designed. Hence, in this study, we utilized AvNAPSA (Asc)^31^ and Rosetta supercharging (Rsc) protocols that have already been deployed in Rosetta software^39^. For a given domain (CBM2a or Cel5A) to achieve the extremes of net charge, positive and negative supercharging protocols were run without a target net charge to ensure that sampling by the software is unbiased by user input. Upon completion of the simulation, the output structure was analyzed in PyMOL to identify whether any amino acids chosen by the software were within 10 Å of the CBM2a binding site or Cel5A active site. In addition, the output structure was analyzed for mutations of helix capping residues and disruption of salt bridges formed between aspartate and arginine residues. Since these amino acid mutations may have deleterious impact on enzyme structure or activity, these residues were included in a resfile and the simulations were re-run with exclusion of these amino acids. Successive iterations of this simulation routine with constraints allowed us to get a better understanding of the net charge range that can be sampled without introducing deleterious mutations. The intent was to design 4 CBM2a mutants and 6 Cel5A mutants, to allow for enough diversity in the overall net charge range sampled. For each target net charge level, mutants from AvNAPSA and Rosetta supercharging protocols were included for the sake of redundancy. Nucleotide sequences for the final designs were codon optimized for *E. coli* and provided to the Joint Genome Institute (Department of Energy) to synthesize designed mutant sequences. Genes for each construct were subcloned between KpnI and XhoI restriction sites in the pET45b(+) expression vector (www.addgene.org/vector-database/2607/). These genes were transformed into T7 SHuffle (New England Biolabs) competent *E. coli* cells and stored as 20% glycerol stocks to be used for enzyme expression described in subsequent sections.

### Small scale protein expression

Glycerol stocks for all CBM2a-Cel5A mutants and wild type full length enzyme were used to inoculate 10 mL of LB media with 100 µg/mL carbenicillin and grown overnight at 37°C, 200 RPM for 16 hours. Overnight cultures were used to make new glycerol stocks for large scale expression, and remaining inoculum was transferred to 200 mL of Studier’s auto-induction medium^40^ (TB+G) with 100 µg/mL carbenicillin. These cultures were incubated for an additional 6 hours at 37°C so that cells could once again reach an exponential growth phase, and protein expression was induced at two different temperatures, first 25 °C for 24 hours, then 16 °C for 20 hours. Cells were pelleted via centrifugation with a Beckman Coulter centrifuge equipped with JA-14 rotor by spinning the entire 200 mL cultures down at 30,100 x g for 10 mins at 4 °C. For lysate characterization, 0.5g of cells were harvested from the main pellet and resuspended with 2.5 mL lysis buffer (20 mM phosphate buffer, 500 mM NaCl, and 20% (v/v) glycerol, pH 7.4), 35 µL protease inhibitor cocktail (1 µM E-64, Sigma Aldrich), and 2.5 µL lysozyme (Sigma Aldrich). Cells were sonicated using a Qsonica Q700 sonicator with 1/8” microtip probe for 1 minute (Amplitude = 20, pulse on time: 5 s, pulse off time: 30 s) on ice to avoid overheating. Insoluble cell debris were pelleted in an Eppendorf 5424 centrifuge with rotor FA-45-24-11 at 15,500 x g for 45 minutes. The soluble lysate supernatant was isolated for biochemical characterization.

### Characterization of soluble cell lysates

Isolated soluble cell lysate activity was characterized using chromogenic substrate pNPC, AFEX pretreated corn stover, cellulose-I, and PASC. Assays with pNPC were described previously^30^. Briefly, 75 µL of 5 mM pNPC prepared in deionized (DI) water was added to 100 µL of soluble cell lysate in 0.2 mL PCR tubes (USA Scientific). All reaction wells and reagents were held on ice to prevent premature reaction before incubation. Reaction wells were then incubated for 30 minutes at 50°C with 200 RPM orbital shaking. After incubating, reaction mixtures were quenched with 25 µL of 0.4 M sodium hydroxide (NaOH) in order to arrest the reaction and increase the pH well above the pKa of 4-nitrophenol. After quenching, 100 µL of reaction supernatant was transferred to a clear flat bottom 96-well microplate (USA Scientific), and endpoint absorbance of pNP (410 nm) was recorded and compared to pNP standards.

Insoluble cellulosic substrates used for hydrolysis assays were prepared as a slurry in deionized (DI) water with 0.2 g/L sodium azide to inhibit microbial contamination. AFEX pretreated corn stover was first milled to 0.5 mm before suspension in DI water at a concentration of 25 g/L, cellulose-I was prepared as a 100 g/L slurry in DI water using Avicel, and PASC was prepared as a 10 g/L slurry with DI water. Hydrolysis assays were conducted by adding 100 µL of substrate slurry (AFEX, cellulose-I, or PASC) and 100 µL of soluble cell lysate to a 0.2 mL 96-well round bottom microplate (Greiner Bio-One). Reaction blanks consisted of cell lysis buffer, protease inhibitor cocktail, and lysozyme were used in place of cell lysate. Microplates were sealed with TPE capmat-96 (Micronic) green plate seals and taped tightly with packing tape on all edges to prevent evaporation. Reaction wells were incubated for four hours at 60 °C in a VWR hybridization oven with end-over-end mixing at 5 RPM. This temperature was chosen based on prior work that found 60°C to be the optimal temperature for *T. fusca* cellulases^41^. Hydrolysis plates were incubated for only four hours as to capture activity prior to 5% total conversion. Concentration of reducing sugars in the soluble hydrolysate supernatant was estimated using dinitrosalicylic acid (DNS) assays^42^ as previously described^43^ and compared to glucose standards.

### Construction of CBM2a-GFP constructs

CBM2a-GFP constructs for select five CBMs used in this study were constructed via Gibson Assembly of the CBM with a green Fluorescent Protein (GFP) insert. In order to keep the architecture of the CBM2a-GFP constructs analogous to their CBM2a-Cel5A counterparts, GFP was fused on the C-terminus of the CBM via the same linker constant throughout all fusion proteins used in this study. Primers were designed to linearize the pET45-b(+) backbone containing both CBM and linker. Although the amino acid sequences are the same for the linker peptide used for all five CBMs, nucleotide sequences differ due to *E. coli* codon optimization mentioned in the previous section, thus five separate pairs of primers were constructed. Insert primers were designed based on GFP from a pEC-GFP-CBM1 DNA template used in previous studies^44^. All primers were synthesized by Integrated DNA Technologies, Inc (IDT) and PCR reactions were conducted following previously published protocols^45^. Remnant wild-type DNA was degraded via DPN1 (New England Biolabs) digestion for two hours at 37°C, and leftover DPN1 enzyme was deactivated by heating the digestion mixture to 80°C for 20 minutes. The remaining PCR products were cleaned via spin columns from an IBI Scientific gel extraction & PCR cleanup kit following manufacturer’s protocols. The final CBM-GFP constructs were assembled using NEBuilder® Hifi DNA Assembly (New England Biolabs) master mix following manufacturers protocols. The cloning mixture was transformed into NEB 5-alpha Competent *E. coli* cells, grown overnight on LB-agar plates at 37°C. Plate colonies were picked at random and used to inoculate 10mL LB media with 100 µg/mL carbenicillin in 15mL culture tubes (VWR). Cultures were once again grown overnight at 37°C, and cells were pelleted via centrifugation at 3,900 RPM in an Eppendorf 5810R centrifuge with rotor S-4-104. Plasmids were extracted using a high-speed miniprep kit (IBI Scientific), sequenced via Sanger sequencing (Azenta), and confirmed sequences were transformed into T7 SHuffle (New England Biolabs) competent *E. coli* cells for large scale expression and purification described in the next section.

### Large scale protein production and purification

Large scale expression of wild type and mutant constructs was done by scaling up protocols from small scale expression. Briefly, 50 mL LB medium and 100 µg/mL carbenicillin was inoculated with glycerol stocks (from small scale cultures) and incubated for 16 hours at 37 °C and 200 RPM. Starter cultures were then transferred to TB+G auto-induction medium and incubated for 6 hours at 37 °C before inducing protein expression at 25 °C for 24 hours then 16°C for 20 hours. Cell pellets were harvested via centrifugation in the same manner described earlier. Entire cell pellets were resuspended with 15mL of cell lysis buffer, 200 µL of protease inhibitor cocktail, and 15 µL of lysozyme per every 3 gram of cell pellets and were vortexed vigorously to evenly suspend the cells. Cells were lysed using a Qsonica Q700 sonicator equipped with a 1/4” microtip for 2.5 minutes (Amplitude = 20, pulse on time: 10s, pulse off time: 30s) on ice. Lysate mixtures were centrifuged at 4°C in an Eppendorf 5810R centrifuge to isolate soluble cell extract. An extra 500 µL of protease inhibitor cocktail was added to the lysates in order to prevent proteolytic cleavage prior to purification. CBM2a-Cel5A proteins were isolated from *E. coli* lysates by immobilized metal affinity chromatography (IMAC) using a BioRad NGC FPLC equipped with a His-trap FF Ni^2+^ − NTA column (GE Healthcare). Columns were regenerated fresh with nickel prior to purification of each sample. Purification was done by first equilibrating the column and system plumbing with start buffer A (100 mM MOPS, 500 mM NaCl, 10 mM imidazole, pH 7.4) at a rate of 5 mL/min for roughly 5 column volumes (25mL). After a achieving a stable baseline via in-line absorbance measured at 280nm, cell lysate was loaded onto into the column at a rate of 2 mL/min. An extra 2 column volumes of buffer A are used to wash the column (bound with his-tagged protein) from impurities until a stable baseline is once again achieved. His-tagged protein is eluted from the column at a rate of 5 mL/min using elution buffer B (100 mM MOPS, 500 mM NaCl, 500 mM imidazole, pH 7.4) and fractions were collected corresponding to A280 peaks. Purity of purified proteins was confirmed via SDS-Page. Size exclusion chromatography using a HiPrep 26/10 desalting column (Cytiva) was done on the NGC system to exchange buffer for storage buffer consisting of 50 mM Mops + 100 mM NaCl pH 7.5 according to the manufacturer’s protocols before long-term storage at −80 °C.

### CBM-GFP pull down binding assay

CBM-GFP pull down binding assays were performed following protocols similar to those described in previous work from our lab^44,46^. All binding assays were performed with six replicates and carried out in 0.2 mL 96-well round bottom microplates (Greiner Bio-One) with crystalline cellulose (Avicel PH-101) prepared as a 100 g/L slurry serving as the cellulose model substrate to screen CBM binding. Protein dilutions were made in 10mM sodium acetate buffer (pH 5.5) for concentrations ranging from 25 to 500 µg/mL. Binding wells consisted of 1 mg total cellulose, bovine serum albumin (BSA) blocking buffer (10 mg/mL BSA + 40 mM sodium acetate pH 5.5), CBM-GFP dilutions, and DI water to top the reaction volume off to 200 µL. Shaken standards and never shaken standards were prepared following a similar composition without any added cellulose. Binding wells and shaken standards were incubated at 25 °C with 5 RPM end over end mixing in a VWR hybridization oven for one hour while never shaken standards were incubated on the lab bench. After incubation, microplates were centrifuged at 3,900 RPM in an Eppendorf 5810R centrifuge for 5 mins to settle cellulose. Results were obtained by carefully transferring 100 µL of supernatant to black 96-well flat bottom plates (VWR), and fluorescence was measured at 480 nm excitation and 512 nm emission with a 495 nm cutoff.

### Cellulose and biomass hydrolysis assays

Purified enzyme activity was screened using both AFEX corn stover and crystalline cellulose-I slurries as described earlier. Reactions were conducted in 0.2 mL round bottom microplates (Greiner Bio-one) at a constant enzyme loading of 120 nmol cellulase / g of substrate. Either 80 µL of AFEX slurry (25 g/L) or 20 µL of cellulose-I slurry (100 g/L) was used so that a total 2 mg of substrate was present in each well. Reactions were composed of either substrate slurry, 50 µL of cellulase dilution (for 120 nmol/g loading), 20 µL of buffer (1 M sodium acetate or sodium phosphate) at a pH within the range of 4.5 – 7.0, and DI water to adjust the final volume to 200 µL DI water. Cellulase dilutions were replaced with 50 µL of DI water for reaction blanks. Microplates were capped with TPE capmat-96 (Micronic) green plate seals and taped tightly with packing tape on all edges to prevent evaporation. Reaction wells were incubated for 24 hours at 60 °C in a VWR hybridization oven with end-over-end mixing at 5 RPM. Microplates were centrifuged to settle solids after incubation so that reducing sugar concentration could be estimated via DNS assay in the manner described earlier.

### Thermal stability assays

Hydrolysis assays were conducted at higher temperatures on both soluble substrate pNPC and insoluble cellulose-I to gauge how supercharging impacted activity at elevated temperatures. Biomass was not used for these experiments as results would be convoluted due to the presence of both soluble xylan and insoluble cellulose in the substrate mixture. Reactions with pNPC were adapted from conducted in 0.2 mL PCR tubes (USA Scientific) and reaction mixtures based on previous protocols^47^ contained 80 µL of 5 mM pNPC, 10 µL cellulase dilution (0.2 nmol of enzyme), and 10 µL of 0.5 M sodium acetate pH 5.5. A pH of 5.5 was chosen based on previous work that found this to be the optimal pH for *T. fusca* cellulases^41^. Reaction tubes were incubated for 30 minutes at a temperature within the range of 55 – 80 °C with orbital shaking at 200 RPMs. Reactions were quenched with 100 µL of 0.1 M NaOH, and reaction mixtures were added to a transparent flat bottom microplate containing 100 µL of DI water. Absorbance of pNP was measured at 410 nm and compared to pNP standards. Assays with cellulose-I were conducted similarly to the methods described in the previous section, except hydrolysis plates were incubated at temperatures within the range of 55 – 80 °C for four hours. Reducing sugar concentration was once again estimated via DNS assay and compared to glucose standards.

## RESULTS AND DISCUSSION

### Computational design of supercharged library

Wild-type CBM2a and wild-type Cel5A carry a net charge of −4 and −2 respectively. Supercharging workflows available in Rosetta software were used to design 4 CBM2a mutants spanning a net charge range of −10 to +8 and 6 Cel5A CD mutants spanning a net charge range of −32 to +44. The four CBM designs cover an even net charge range compared to the wild type CBM, with two designs negatively supercharged (D1 (net charge: −10) & D2 (net charge: −8)) and two designs positively charged (D3 (net charge: +6) & D4 (net charge: +8)). The net charge increases with design number from D1 (most negative) to D4 (most positive) as shown in **Figure 1**. Similarly, the six Cel5A designs cover a net charge range of −32 to +44 with two designs negatively supercharged (D1 (net charge: −32) & D2 (net charge: −29)) and four designs positively supercharged (D3 (net charge: +11), D4 (net charge: +14), D5 (net charge: +41) and D6 (net charge: +44)). The mutations necessary to create each of these designs are summarized in **Supplementary Table T1** and **Supplementary Table T2** while the amino acid sequences for wild-type CBM2a and wild-type Cel5A can be located in the supplementary sequences excel file.

**Figure 1.**
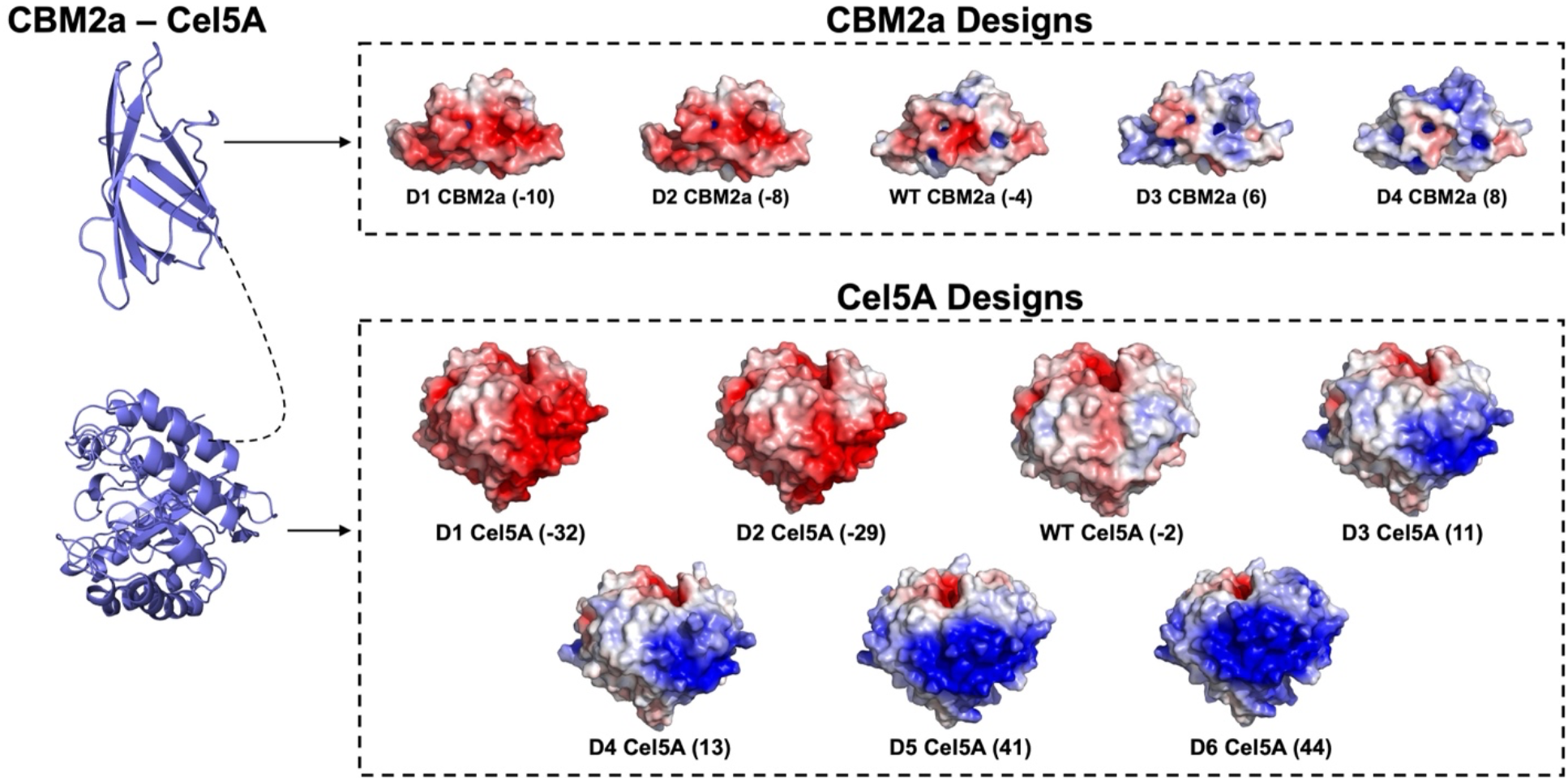
Computational design of CBM2a and Cel5A mutants and library construction. Rosetta macromolecular software was used to identify surface amino acid residues on the surface of either CBM2a or Cel5a for mutation to positively charged (K, R) or negatively charged (D, E) amino acids. Each domain was mutated individually then one of five CBM designs was fused with one of seven Cel5A CD designs via a flexible linker peptide creating a total possible library size of 35 mutants. Net charges for each design or wildtype domain (WT) are indicated in parenthesis and were estimated by the total number of charged amino acid residues. Electrostatic potential maps ranging from −5 kT/e (red) to +5 kT/e (blue) were generated using Adaptive Poisson – Boltzman Solver (APBS) plugin in PyMOL.

The net charge range chosen for each domain and the granularity of net charge sampling were decided based on the following considerations: 1. Running unconstrained simulations (without a target net charge) allows one to obtain the maximum target net charge that does not cause significant structural perturbation of the protein 2. Rosetta supercharging algorithm may predict different mutations from the AvNAPSA algorithm and hence the construction of a supercharging library with sequence diversity should feature both approaches. Hence, upon deciding the target net charge range, an attempt was made to obtain a design each with Rosetta and AvNAPSA, that possess similar net charges. For instance, D1 and D2 CBM2a are obtained using different approaches but possess net charges in close proximity (−10 and −8 respectively).

Altogether, including the wild type CBM and CD, there are 5 CBM constructs and 7 CD constructs. Each CBM is fused to a CD construct via flexible linker that is constant for all constructs bringing the total library size to 35 mutants. Two mutant sequences were unable to be synthesized by the JGI (WT CBM2a – D5 Cel5A and D4 CBM2a – D4 Cel5A) reducing the total library size down to 33 mutants. Constructs were received as glycerol stocks with DNA already inserted into pET45b(+), and construct validation was performed to ensure correct sequence identity by sequencing a pool of random mutants picked via random number generator. Constructs that were expressed on a large scale were additionally sequenced to confirm their identity prior to further characterization.

### Screening of entire library based on soluble cell lysate activity

In order to understand how well the supercharged constructs expressed as well as characterize activity on different cellulosic substrates, all 32 mutants and wild-type CBM2a-Cel5A enzymes were expressed in *E. coli* and the resulting soluble cell lysates were used for biochemical characterization. It is important to note that enzyme loading is not fixed, thus differences in activity observed may arise from a change in catalytic turnover, or from differences in expression levels. Assays using soluble chromogenic para-nitrophenyl cellobiose provide rough insight on whether or not the enzymes are expressing and if they are active (**Figure 2A**). pNPC assays show low overall activity for most constructs with the exception of three constructs containing a mutated CBM and wildtype CD. Of these, the D2 CBM2a WT Cel5A construct containing a negatively supercharged CBM and D3 CBM2a – WT Cel5A construct carrying a positively supercharged CBM stand out as being more active than the wildtype enzyme (Construct 1). Interactions with soluble pNPC occur with only the Cel5A active site^48^, thus mutations to the CBM aren’t expected to produce drastic activity changes when hydrolyzing pNPC. The increase in pNP hydrolysis for the two CBM mutants (D2 & D3; Constructs 13&20) containing wildtype Cel5A may be a result of differences in expression levels, or improvements in solubility at the pH tested occurring as a result of changing the enzyme’s pI. Most other enzymes showed either zero or low activity on pNPC which at first glance implies that these enzymes are either not expressing, or not active. Results on insoluble substrates like biomass and cellulose-I indicate this is not the case. One potential cause for the low activity on pNPC may be due to a decrease in thermostability for many of the constructs, especially those with mutated Cel5A catalytic domains. Enzyme binding to substrate has been shown in the past to help stabilize the enzymes at elevated temperatures^49^. Thus, without insoluble substrate present to form a stable enzyme-substrate complex, many of the CBM2a-Cel5A variants are subject to thermal denaturation and subsequent unfolding and precipitation resulting in low activity on pNPC. Alternatively, the charged residues might be allosterically interacting with pNPC to impact activity. All 32 mutants exhibit much higher activity in comparison to pNPC assays on insoluble biomass (**Figure 2B**), cellulose I (**Figure 2C**), and PASC (**Supplementary Figure 1**). Several mutants stand out as exhibiting higher activity on cellulosic substrates compared to the wildtype enzyme. In general, the greatest activity is seen when one mutant domain (either CBM or CD) is coupled with a wildtype domain. This may once again be related to expression and protein folding, with highly mutated and drastically charged species expressing poorly or misfolding compared to other mutants. From this list of constructs with one mutated domain, all four CBM mutants (D1-D4 CBM2a −WT Cel5A) and three CD mutants (WT CBM2a − D2-D4 Cel5A) have similar or greater activity on two or more insoluble substrates compared to the wildtype enzyme. Several combinatorial constructs that contain two mutated domains also showed higher activity, with most of these mutants containing the D2, D3, or D4 CD that exhibited higher standalone activity. It is important to note that improvements to activity do not appear to be additive. For example, on pretreated biomass, the D3 CBM (D3 CBM2a – WT Cel5A) and D3 CD mutant (WT CBM2a – D3 Cel5A) are the two most active constructs. Interestingly though, the combination of these two domains together (Construct 17, D3 CBM2a – D3 Cel5A) only ever produces half the activity of the wildtype enzyme at best and has a more than two-fold decrease compared to the D3 constructs with only one mutant domain. It does appear from lysate activities that supercharging only one domain is more effective at increasing hydrolysis yield on biomass and cellulose substrates, with mutations of the CBM being more effective at improving activity.

**Figure 2.**
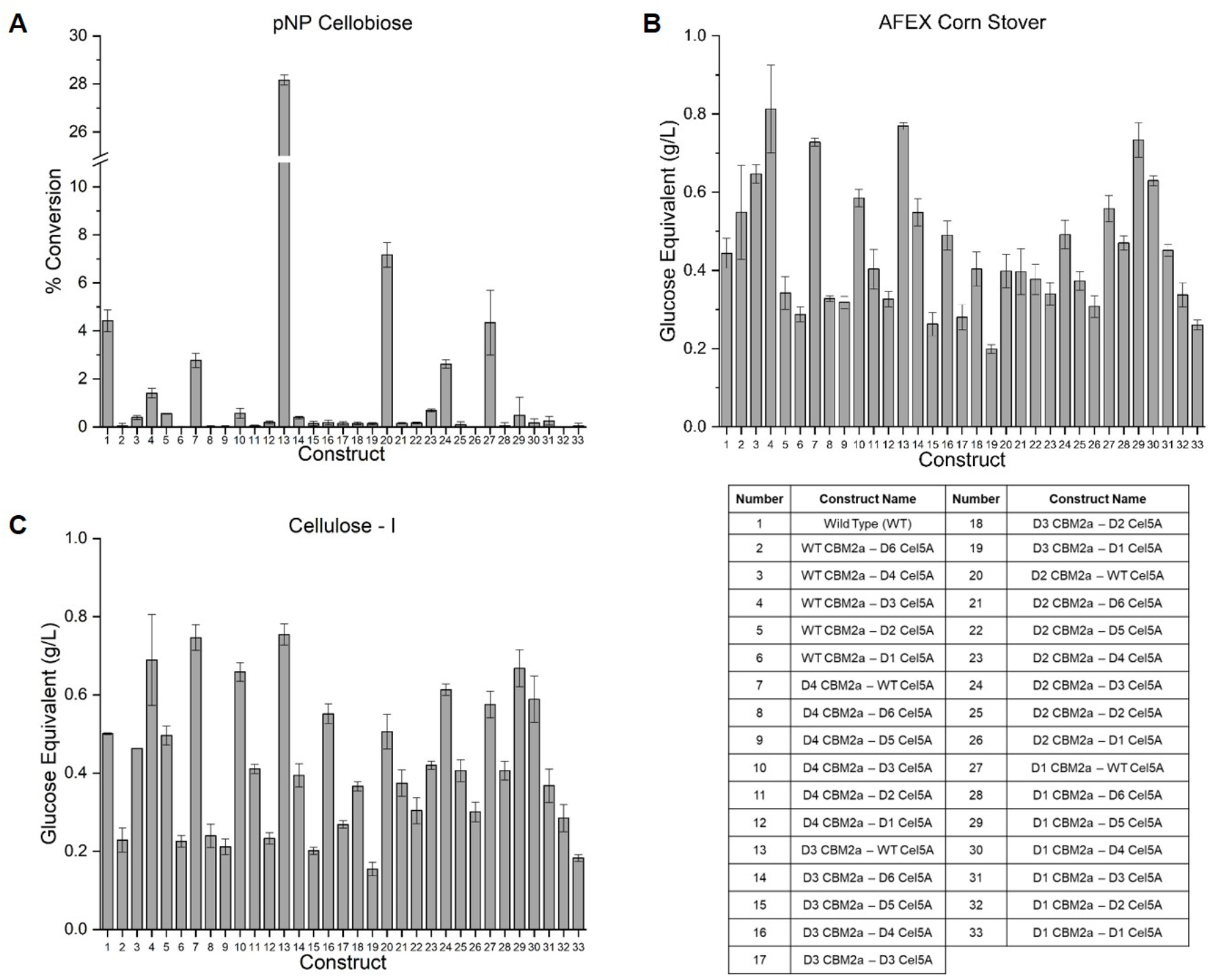
Screening soluble cell lysates for entire library identifies several CBM2a – Cel5A mutant constructs with higher catalytic activity than wildtype full length enzyme. All 33 constructs depicted were expressed as 200mL auto-induction cultures and pellets harvested via centrifugation were sonicated in buffer containing 20 mM sodium phosphate pH 7.4, 500 mL sodium chloride, and 20% (v/v) glycerol. (A) Para-nitrophenyl cellobiose (pNPC) was used to characterize soluble substrate activity by incubating 100 µL of isolated soluble cell lysate with 5mM pNPC for 30 mins at 50°C before quenching with sodium hydroxide. Absorbance at 410nm was used to estimate the percent of pNPC converted to yellow-colored paranitrophenol (pNP) by comparing to pNP standards. (B) Amonia fiber expansion pretreated (AFEX) cornstover was prepared as a 25 g/L slurry in DI water and 100 µL of slurry was incubated with 100 µL of soluble cell lysate for 6 hours at 60 °C. Hydrolysate supernatant was isolated via centrifugation and reducing sugar concentration was estimated using DNS assay and compared to glucose standards. (C) Crystalline Cellulose – I was prepared from Avicel PH – 101 as a 100 g/L slurry and incubated with 100µL soluble cell lysate for 6 hours at 60 °C. Hydrolysate supernatant was obtained via centrifugation, and DNS assay was used to estimate reducing sugar concentration in the hydrolysis supernatant. All data points represent the average of four technical replicates and error bars represent one standard deviation.

When comparing groups of mutants that contain the same CBM but a different CD mutant another interesting trend is observed. For some of these “sub-families”, there is a near unimodal distribution of activity for each member. This trend is readily visible for the WT CBM sub-family (Constructs 2 − 6) and D1 CBM sub-family (Constructs 28 − 33). In these groups of constructs, activity increases from one design to another until a clear peak is reached, then activity will steadily decrease for the subsequent constructs afterwards. Therefore, there appears to be a “sweet spot” or optimal net charge corresponding to each design where the activity is maximized. This phenomena resembles that of a Sabatier optimum^32,33^ where in this case a specific net charge likely modulates binding affinity to insoluble substrates so that a specific net charge provides an intermediary binding strength in order to maximize catalytic turnover. It is likely that this optimal charge will be different for different substrates, but this is not observed through lysate screening most likely due to assay conditions. This trend loosely holds for each sub-family, where those outliers may exist because of other factors such as low expression, low solubility, low thermostability, etc. Absolute activity of all 32 mutants and wildtype CBM2a-Cel5A is summarized in **Supplementary Table T3** with constructs exhibiting higher activities on two or more substrates highlighted in green. These results also include a T7 Shuffle empty vector control to ensure there was no background catalytic activity being measured on any substrate from the T7 background lysate.

### Positively charged CBMs bind cellulose with a higher affinity

Based on our previous work^30^ it is hypothesized that electrostatic interactions between supercharged CBMs and crystalline cellulose can significantly alter CBM binding and resulting catalytic activity of the full length enzymes. To elucidate the impact CBM net charge has on cellulose binding, fluorescence-based pull-down binding assays were performed for three supercharged constructs (D2-D4) and the native CBM. The most negatively charged CBM (D1 CBM2a) was omitted due to difficulties in expressing and purifying the negative GFP tagged CBM in *E. coli*. Binding data on crystalline cellulose-I for all four constructs was fit to a one-site Langmuir isotherm model (R^2^ ≥ 0.95). The maximum number of binding sites (N_max_) on cellulose, binding dissociation constant (K_d_), and partition coefficient (N_max_/K_d_) were estimated from the fits and are listed in **Table 1**. Recent work from our group has identified that GFP binding to cellulose is insignificant with roughly two orders of magnitude lower available binding sites than CBM tagged GFP^50^, and thus the contribution of GFP binding to cellulose has been assumed to be negligible.

**Table 1.** Binding parameters for wild-type and supercharged CBMs from Langmuir one-site isotherm. CBM charge at pH 5.5 refers to the charge of only the CBM. Binding parameters Nmax and Kd, along with their respective standard deviation was obtained by fitting pull-down binding assay data to a Langmuir one-site model in Origin software. Partition coefficient (Nmax/Kd) was obtained by dividing the model parameters in columns three and four.

| Construct | CBM Charge<br>(pH 5.5) | $N_{\max}$<br>($\mu\text{mol/g}$<br>Cellulose) | $K_d$ ($\mu\text{M}$) | $N_{\max}/K_d$<br>(L/g Cellulose) |
| --- | --- | --- | --- | --- |
| D4 CBM2a | 8 | $2.86 \pm 0.63$ | $0.33 \pm 0.12$ | $8.64 \pm 0.41$ |
| D3 CBM2a | 6.5 | $3.11 \pm 0.57$ | $0.51 \pm 0.15$ | $6.11 \pm 0.35$ |
| WT CBM2a | -3.5 | $2.08 \pm 0.12$ | $1.11 \pm 0.15$ | $1.87 \pm 0.15$ |
| D2 CBM2a | -7.5 | $2.49 \pm 0.65$ | $2.96 \pm 1.01$ | $0.84 \pm 0.43$ |

The charge of each CBM (excluding GFP) at pH 5.5 was estimated with an online charge calculator (protcalc.sourceforge.net/) using the primary sequences of each construct. These charges were correlated to binding parameters for each construct (**Figure 3**) to elucidate how charge differences impact cellulose binding. There is no apparent difference in N_max_ for the constructs tested (**Figure 3A**) suggesting that supercharging has not altered the amount of available binding sites accessed by the CBM. However, supercharging has noticeable altered the binding dissociation constant. **Figure 3B** suggests that binding affinity (approximated as the inverse of K_d_) increases with CBM net charge, with the most positively charged CBM construct (D4 CBM2a) having a dissociation constant more than 3-fold lower (or 3-fold higher association constant) than the wild-type CBM. Partition coefficients for each CBM relating the amount of enzyme bound to cellulose to free enzyme in solution can be found from the slope of the linear portion of the binding curves. These linear portions have been plotted on the same axes (**Figure 3C**) to visualize how charge impacts the partition coefficient. Once again there is a direct correlation to the charge of each CBM, with the most negative CBM tested (D2 CBM2a) exhibiting the lowest partition coefficient, and the most positive (D4 CBM2a) showing the highest. These results imply that increasing positive charge on the CBM does improve binding to cellulose clearly identified by decreased dissociation constants and increased partition coefficients. This effect certainly contributes to the activity improvements observed in lysate screening for some of the positively supercharged CBM constructs, but it is not yet clear what the limit to this effect is where strong binding leads to dissociation limitations, and thus decreased activity.

**Figure 3.**
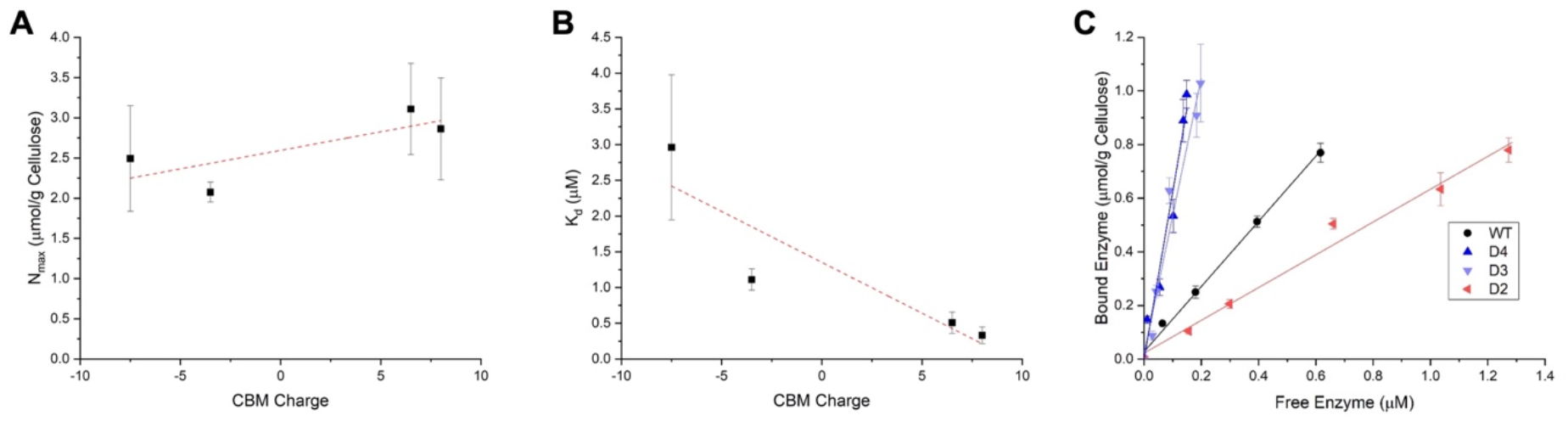
Supercharging does not impact Nmax, but significantly alters binding affinity and partition coefficient on cellulose-I. GFP tagged constructs were expressed as 1L auto-induction cultures, and N-terminus his-tagged enzymes were purified from *E. coli* lysate by immobilized metal affinity chromatography. Pull down binding assays were performed with a total 1 mg insoluble cellulose, 2.5 mg/mL BSA, 10 mM NaOAc pH 5.5, and protein dilutions ranging from 25 – 500 µg/mL made using NaOAc buffer. Cellulose was replaced with DI water for shaken and never shaken standards prepared alongside binding wells. All plates were incubated for one hour at 25 °C with binding wells and shaken standards being mixed end-over-end at 5 RPM, and never shaken standards kept on the lab bench. After incubation all plates were centrifuged at 3,900 RPM, and 100 µL of supernatant was transferred to opaque flat bottom microplates to measure fluorescence at 480 excitation and 512 emissions with 495 nm cutoff. Full scale binding curves presented in the supplementary information were constructed using Origin plotting software and data was fit to a one-site Langmuir isotherm model. (A) Maximum number of binding sites (Nmax) on cellulose resulting from one-site model fit for all four CBM-GFP constructs plotted as a function of the corresponding CBM charge. CBM charge refers to the charge of the binding module only and was estimated using the primary amino acid sequence for each CBM using an online protein charge calculator (https://protcalc.sourceforge.net/). (B) Binding dissociation constant (Kd) found from one-site model fits for all four CBM constructs plotted as a function of CBM charge. (C) Linear portion of binding curves for all four constructs tested. The slope of these plots corresponds to the partition coefficient for each CBM construct which can be defined as (Nmax/Kd). All data reported represents the average of six technical replicates, and error bars represent standard deviation from the mean.

### Supercharging both CBM and CD shifts pH optimum

The isoelectric point (pI) of proteins describing the pH where proteins have zero net charge is dictated by ionizable groups within the side chains of the primary amino acid sequence. Altering protein net charge by introducing charged amino acids has been shown to alter solubility, and can potentially shift the pH where maximum activity is observed^51^. The process of supercharging significantly shifts the pI of CBM2a-Cel5A mutants through the manipulation of charged amino acid residues (D, E, R, K) present on the protein’s surface. To understand how this has impacted pH dependence, all four CBM mutants (supercharged CBM, wildtype CD), and the three CD mutants (wildtype CBM, mutant CD) that showed activity improvements through lysate screening (D2, D3, D4) were expressed and purified, and their activities characterized on AFEX corn stover and cellulose-I and compared to wild type CBM2a-Cel5A. The wildtype full length enzyme is expected to have an optimal pH of 5.5 based on previous work utilizing pretreated biomass and crystalline cellulose substrates^41^, but activity on biomass and cellulose-I (**Figure 4**) showed that this optimal pH is actually closer to pH 6.0. When comparing CBM mutant activity in the pH range of 4.5 – 7.0 on biomass, (**Figure 4A**) differences in optimal pH can be observed. For one, the negative D2 CBM mutant clearly shows greatest activity at pH 5.5, where it is nearly two times more active when compared to the wildtype enzyme at its pH optimum. Interestingly, the D2 CBM mutant is much more active than all other mutants and wildtype enzyme at the more acidic range of pH’s tested. The other negative CBM mutant (D1 CBM) does not exhibit the same behavior and shares the same optimal pH as the wildtype enzyme where it is similar, if not slightly more active, but otherwise this mutant does not provide the improvements that D2 does on biomass. The positively supercharged CBMs (D3 and D4) display peak activity past a pH of 6.0, and in this pH range are also more than two times more active compared to the wildtype enzyme. These activity improvements are likely related to the net charge of the positive mutants. Substrates containing lignin like corn stover will non-productively bind to enzymes, and this effect is stronger for positively charged enzymes^29^. As the pH increases and ionizable groups are deprotonated, the net charge of these constructs will become less positive, reducing the effect of non-productive binding towards lignin. Within this range, both mutants show improved activity on biomass, and the D3 CBM mutant retains this activity up to a neutral pH. On the other hand, the three CD mutants do not show nearly the same improvements on biomass (**Figure 4B**). Of the three, the negative D2 CD mutant is most active at pH 5.5 where its activity is significantly decreased compared to wildtype. The two positive CD mutants tested (D3 and D4) respond to changes in pH in a similar fashion compared to the positive CBM mutants, but at best are only equal in activity to the wildtype enzyme near neutral pH.

**Figure 4.**
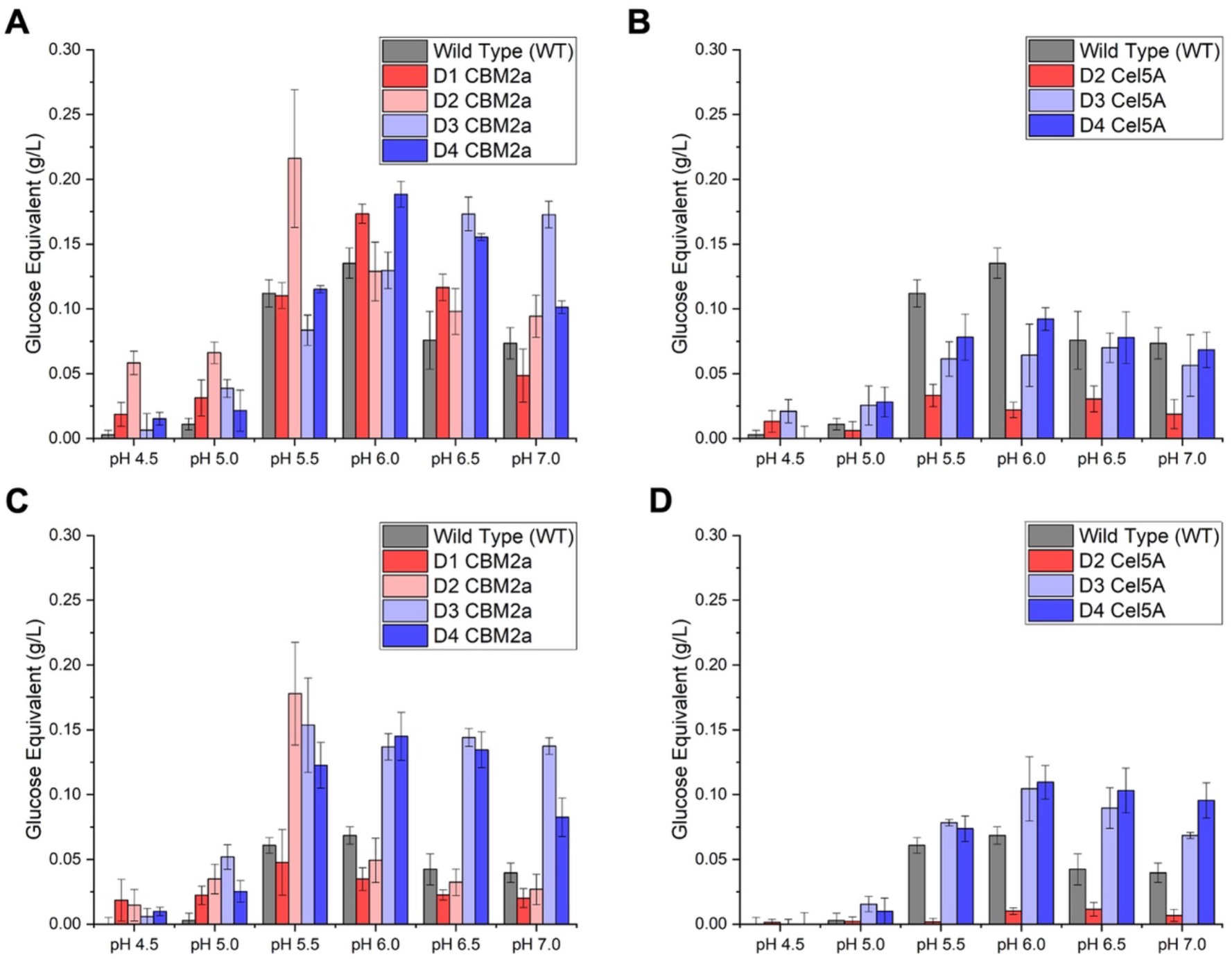
Catalytic activity of purified single mutant domains is highly substrate and pH dependent with large activity improvements observed in positively supercharged domains on crystalline Cellulose – I. Enzymes were expressed as 1L auto-induction cultures, and N-terminus his-tagged enzymes were purified from *E. coli* lysate by immobilized metal affinity chromatography. Purified enzyme assays conducted with insoluble substrate slurries consisted of 120 nmol enzyme per gram of substrate with a total 2mg of substrate per reaction mixture. Enzyme assays were conducted in buffers ranging in pH from 4.5 – 7.0 and were incubated 24 hours at 60°C. All dilutions were made in deionized water and the minimal amount of salt was added in order to observe full effects of net charge unabated by charge screening. Reducing sugar equivalents were estimated via DNS assay and compared to glucose standards. All mutants shown are color coded from most negative (red) to most positive (blue). (A) Glucose equivalents released after 24-hour AFEX cornstover hydrolysis for wildtype full length enzyme and four CBM2a mutant constructs. All five enzymes tested are full length containing the native wildtype Cel5A. (B) Glucose equivalents released after 24-hour AFEX cornstover hydrolysis for wildtype enzyme and three Cel5A CD mutant constructs. All four enzymes tested are full length containing the native wildtype CBM2a domain. (C) Glucose equivalents released after 24-hour crystalline cellulose – I hydrolysis for the wildtype enzyme and four CBM2a mutant constructs. All five enzymes tested are full length containing the native wildtype Cel5A catalytic domain. (D) Glucose equivalents released after 24-hour crystalline cellulose-I hydrolysis for wildtype enzyme and three Cel5A CD mutants. All four enzymes tested are full length containing the native wildtype CBM2a binding module. All data reported represents the average of four technical replicates, and error bars represent standard deviation from the mean.

Activity screened at different pH on crystalline cellulose-I (**Figure 4C**) once again depicts that the optimal pH for wildtype CBM2a-Cel5A is closer to pH 6.0 and even at this optimum the wildtype enzyme is less active on cellulose-I compared to biomass. This low activity seems to have been significantly improved by positively supercharging the CBM domain. Both positive CBM mutants are around 2-fold more active on cellulose-I than the wildtype enzyme past pH 5.0, and the D3 CBM mutant retains this activity up to a neutral pH. A similar trend is observed for the positively supercharged CD mutants as well (**Figure 4D**), but activity improvements are not as pronounced as the two positively supercharged CBMs. Similar to previous results observed from Whithead et al. (2017)^30^, negatively supercharging both CBM and CD significantly reduced hydrolytic activity on crystalline cellulose. Both D1 CBM mutant and D2 CD mutant showed less activity than the wildtype enzyme, with the D2 CD mutant being virtually inactive on cellulose-I at every pH. These interactions, both favorable and unfavorable, can be attributed to electrostatic interactions between charged enzyme and the negatively charged cellulose substrate. For the positively supercharged mutants, supercharging increases activity on cellulose due to favorable coulombic attraction with the negative substrate surface, whereas negatively supercharged mutants exhibit lower activity due to poorer binding to cellulose arising from electrostatic repulsion with the cellulose surface. These trends can be manipulated by the addition of salt (**Supplementary Figure 4A**) where adding NaCl to screen charges decreases activity on cellulose for positive mutants near their pH optimum and improves activity for negative mutants near their optimum. The D2 CBM mutant is an exception to this trend and behaves as an outlier at pH 5.5. At every other pH tested, the D2 CBM mutant showed similar or lower activity than wildtype, but a sharp peak in activity occurs at pH 5.5 where it is nearly 2.5-fold more active than the wildtype. The cause for this improvement only at the optimal pH is not totally clear. There still appear to be unfavorable interactions occurring between the negatively charged enzyme and cellulose substrate as this activity is even further improved by screening charges with the addition of salt to nearly a 5-fold increase in activity compared to wildtype (**Supplementary Figure 4B**). One hypothesis is that the negative charges introduced on the D2 CBM mutant, while detrimental to adsorption, increase enzyme desorption or reduce non-productive binding to cellulose^44^. Below the pH optimum desorption is less favorable due to a low negative charge, and past the pH optimum adsorption is significantly hindered by high negative charges. The D2 CBM mutant differs from the D1 mutant by only two mutations that are adjacent to two planar aromatic residues. These mutations may further limit adsorption to cellulose by electrostatic repulsion, explaining why the D1 CBM mutant does not experience similar activity improvements. Homology modelling of the supercharged CBM constructs (**Supplementary Figure 5**) indicates changes in tertiary structure for each CBM mutant, along with the orientation of their planar aromatic amino acid residues that dictate binding to cellulose. These changes may have resulted in changes to the CBM structure-function, but this is beyond the scope of this current work.

### Peak catalytic activity observed is correlated to net charge

Changing the solution pH in which the enzymes are characterized subsequently changes the enzyme’s net charge. Using the amino acid sequence for each construct, the net charge was calculated using online tools at each pH value tested in the previous section. These results were correlated to the measured enzyme activity to understand the impact that net charge has on activity on both biomass and cellulose-I for the wildtype, CBM mutants, and CD mutants. **Figure 5** depicts that on both substrates there appears to be a net charge where optimal activity is observed, and these peaks are different on either substrate. In the case of AFEX corn stover (**Figure 5A**), the highest activity occurs at a net charge of around −10, with majority of the constructs screened in the range from 0 to −20 showing moderate to high activity. There seems to be a good correlation between different constructs, with both wildtype and CBM mutants peaking at around the same net charge, and both CBM and CD mutants have similar activity at similar net charges. It is interesting to note that the three purified CD constructs never showed much activity on biomass, and none of these constructs were near the net charge peak when screened on biomass. Results on cellulose-I (**Figure 5B**) show a much tighter relationship with a near unimodal distribution corresponding to a peak in activity around a slight positive charge of 5. Once again there is good correlation across all three groups of constructs, with the wildtype nearly matching activity of CBM mutants at negative net charges. Results on both substrates report that net charge is a good predictor of enzyme activity on different substrates, but different substrates require different optimal charges. For substrates containing lignin such as lignocellulose biomass (e.g., corn stover), it is clearly beneficial to have negatively charged enzymes in order to ease lignin inhibition. On the other hand, for cellulosic substrates that contain no lignin and slight negative surface charges (cellulose-I), it is more favorable to have slight negative charges to improve adsorption through coulombic attraction. However, net charge does not appear to be the sole predictor of enzyme activity. When the wildtype enzyme reaches the peak net charge (+5) on cellulose I, it is nearly inactive. Similarly, although the CD mutant activity peaks in this same charge range, they are still not as active as the CBM mutant. Therefore, while altering net charge is effective at modifying activity and optimizing performance for different substrates, it is not the only factor controlling changes in catalytic activity.

**Figure 5.**
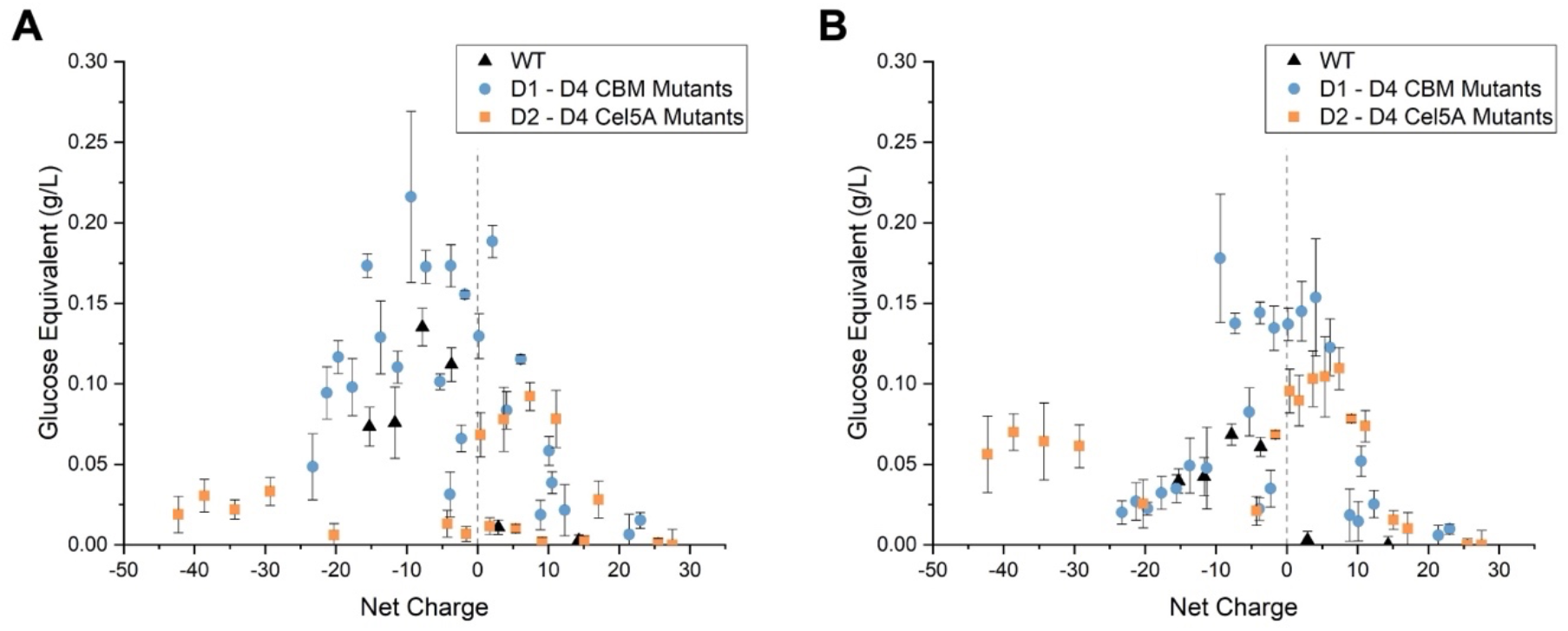
Enhanced catalytic activity correlates to overall net charge in a substrate dependent relationship. Enzyme activity from pH screening assays was adapted and plotted as a function of full-length net charge. Enzyme net charge was calculated at each pH tested from Figure 3 based on full length enzyme sequences. The grey dashed line designates the origin where net charge is zero. (A) Glucose equivalents from AFEX hydrolysis correlated to full length enzyme net charge for wildtype full length enzyme (black triangles), all four CBM mutans (blue circles; Fig 3A), and three Cel5A mutants (orange square; Fig3B). (B) Glucose equivalents from cellulose-I hydrolysis correlated to full length enzyme net charge for wildtype full length enzyme (black triangles), all four CBM mutans (blue circles; Fig 3C), and three Cel5A mutants (orange square; Fig3D). All data points are averages of four technical replicates and error bars represent standard deviation from the mean. Net charge based on the enzyme sequences were estimated using an online charge calculator (https://protcalc.sourceforge.net/).

### Positively supercharged CBMs show increased optimal temperature on cellulose

Assays run on pNPC and cellulose-I were incubated at elevated temperatures to understand how supercharging impacted enzyme stability (**Figure 6**). AFEX corn stover was omitted from these assays as results would be convoluted due to the presence of soluble hemicellulose in the biomass matrix. Aside from the negatively supercharged D2 CD mutant, all enzymes exhibit optimal activity on pNPC at 65 ℃ (**Figure 6A**). This is a drastic difference from the temperature used for lysate screening on pNPC (50℃) that was chosen based on previous protocols analyzing cellulase activity on soluble substrates^52^. At this temperature optimum, the wildtype, D2 CBM construct, and D3 CD construct are roughly equal in activity confirmed by student’s T-test p-values (**Supplementary Table T6**). There does not appear to be any correlation between engineering of the CBM or CD, or any preference to positive or negative supercharging. At 70℃ it is evident that there is not much of an improvement to thermostability, as the wildtype enzyme still retains 90% activity at this point while the mutants lose more than 20% of their optimal activity. However, results on cellulose-I (**Figure 6B**) demonstrate that the two positively supercharged CBMs are more tolerant to high temperatures in the presence of insoluble substrate. Both D3 and D4 CBM mutants show a five-degree higher optimal temperature (65℃) than all other constructs including the wildtype where cellulose hydrolysis is increased 1.5-fold. Additionally, the D3 CBM mutant has nearly the same activity at 70℃ as the wildtype does at its temperature optimum (60℃) indicating a nearly 10-degree increase in thermostability in the presence of substrate. Both mutants remain more active than the wildtype up to 75℃ further displaying their stability. To further examine the impact of this improved thermostability, both CBM mutants and wildtype enzyme were incubated with cellulose at 65℃ for longer time frames (**Figure 6C**). Some variations in later timepoints can be observed, likely due to issues related to evaporation losses when incubating at longer times and higher temperatures. Both the wildtype enzyme and D4 CBM mutant seems to nearly level off after 6 hrs of hydrolysis with a 2-fold difference in activity between the two constructs and wildtype enzyme. However, the D3 CBM mutant seems to remain active past this point, and after 72 hours, it was shown to release 2.5-fold more glucose than the wildtype enzyme by nature of its improved thermostability at the elevated temperature optimum. Substrate stabilization of enzymes at higher temperatures has been recorded in the past^49,53,54^, and in this scenario it can be hypothesized that the favorable surface charged interactions between the negative substrate and positive binding modules seems to even further increase this stabilization resulting in improved thermotolerance, and a resulting increase in turnover due to the higher temperature.

**Figure 6.**
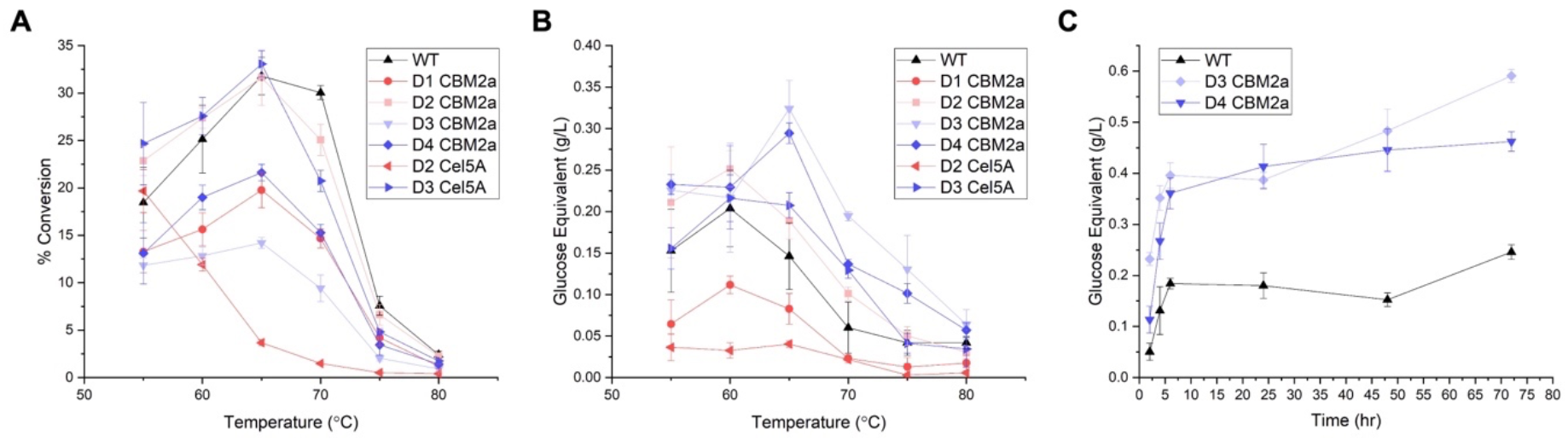
Supercharging enhances optimal hydrolysis temperature for D3 and D4 CBM2a constructs (WT Cel5A) in the presence of cellulosic substrates. (A) Percent conversion of pNPC to pNP as a function of incubation for wildtype enzyme, all four CBM2a mutants (WT Cel5A), one negative, and one positive Cel5A CD mutant (WT CBM2a). Hydrolysis reactions consisted 5mM pNPC stock hydrolyzed by a total of 0.2nmol of enzyme. Reaction mixtures were incubated for 30 mins at the temperature designated on the x-axis, and reactions were quenched with sodium hydroxide after incubation. The concentration of yellow colored pNP released was estimated using absorbance values measured at 410 nm and comparison to pNP standards ranging in concentration from 0 to 5 mM. (B) Glucose equivalents as a function of incubation temperature yielded after hydrolysis of cellulose-I with wildtype enzyme, all four CBM2a mutants (WT Cel5A), one negative, and one positive Cel5A CD mutant (WT CBM2a). A total of 4mg of cellulose-I prepared as a 100 g/L slurry from Avicel PH-101 was incubated with a total enzyme loading of 120nmol/g for four hours at the temperatures indicated on the x-axis. Reducing sugar concentration in the soluble hydrolysate was estimated via DNS reducing sugar assay and glucose equivalents quantified from glucose standards. (C) Based on the data reported from (B), the two best performing mutants (D3 CBM2a – WT Cel5A and D4 CBM2a – WT Cel5A) which exhibited a higher overall optimal temperature were examined at this new optimum. A total of 4mg of cellulose-I was incubated at the inflated temperature optimum (65 °C) with 120 nmol enzyme per gram of substrate for longer time periods up to 72 hours. Data was recorded by removing hydrolysis reactions from their incubators and holding at −20 °C to arrest the reaction at the time points designated on the x-axis. Reducing sugar concentration was estimated via DNS reducing sugar assay and compared to glucose standards. All data reported represents the average of four technical replicates and error bars represent standard deviation from the mean.

### Combing improved supercharged domains does not lead to additive improvements

Biochemical assays with purified enzymes identify three CBM designs (D2, D3, D4) and two CD designs (D3, D4) that, when coupled with a wildtype domain, showed either improved thermostability or higher catalytic activity than the wildtype full length enzyme. Of the six possible chimeras produced by combining an improved CBM and CD, two constructs showed lower activities in lysate screening (D3 CBM2a – D3 Cel5A & D2 CBM2a – D4 Cel5A), and one was not synthesized (D4 CBM2a – D4 Cel5A). The remaining constructs (D2 CBM2a – D3 Cel5A, D3 CBM3a – D4 Cel5A, D4 CBM2a – D3 Cel5A) were expressed, purified, and characterized in a similar manner as the CBM and CD mutant constructs described in the previous sections. In order to deconvolute the impact of combining two supercharged domains, combinatorial mutant activity was screened at different pH on pretreated biomass (**Figure 7 A-C**) and crystalline cellulose (**Figure 7 D-F**). In nearly every case, activity of the combinatorial mutants is constrained by the activity of the individual CBM or CD mutant; there is no additive increase in activity when combing two mutated domains. In scenarios were the individual CBM and CD mutants shared similar optimal pH, the combinatorial mutant had similar activity to either one of the individual mutants. This is evident for both D4 CBM2a – D3 Cel5A and D3 CBM2a – D4 Cel5A which both contain two positively supercharged domains. Screening individual CBM mutants and CD mutants in the previous section showed greater improvements to overall activity when only the CBM was mutated with upwards of a twofold difference in activity between CBM mutants and CD mutants on biomass. This low biomass activity for the mutated Cel5A domains significantly dampens combinatorial mutant activity on pretreated biomass, eliminating the activity improvements observed when the CBM mutant alone was mutated. On crystalline cellulose, both positively supercharged combinatorial mutants show activities either between that of the individual mutants, or below them. Once again, it appears that supercharging only the CBM provides greater contributions to improving catalytic performance. At pH values past 5.5 all positively supercharged constructs are more active than the wildtype enzyme, but once again constructs with only a positively supercharged CBM (D3 and D4) and a wildtype Cel5A CD still remain the most active across a span of solution conditions.

**Figure 7.**
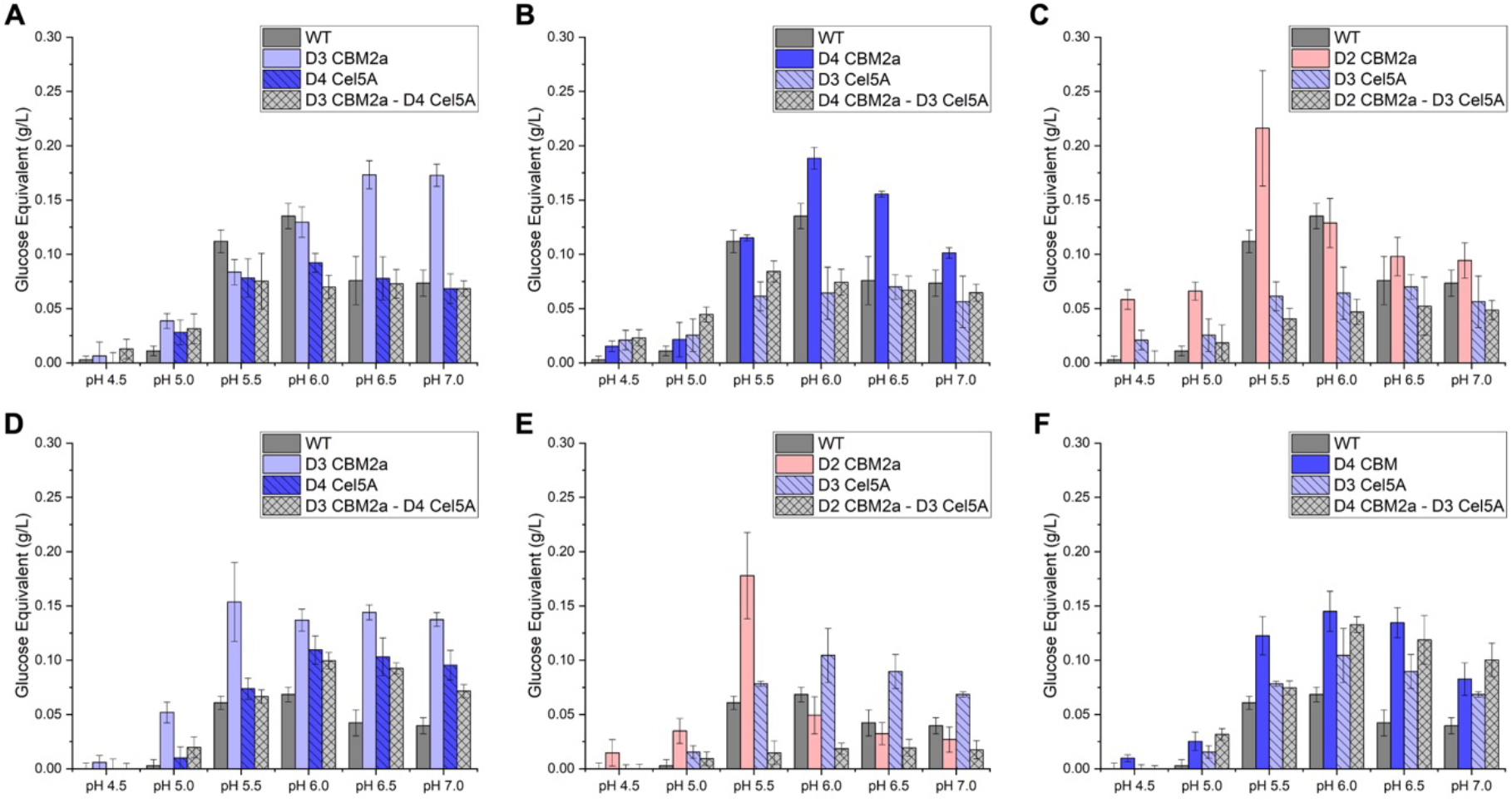
Combining two mutated domains does not lead to an additive improvement in catalytic activity. Combinatorial mutants comprising of the best performing CBM2a or Cel5A designs were expressed as 1L auto-induction cultures, and N-terminus his-tagged enzymes were purified from *E. coli* lysate by immobilized metal affinity chromatography. Purified enzyme assays conducted with insoluble substrate slurries consisted of 120 nmol enzyme per gram of substrate with a total 2mg of substrate per reaction mixture. Enzyme assays were conducted in buffers ranging in pH from 4.5 – 7.0 and were incubated 24 hours at 60°C. All dilutions were made in deionized water and the minimal amount of salt was added in order to observe full effects of net charge unabated by charge screening. Reducing sugar equivalents were estimated via DNS assay and compared to glucose standards. Combinatorial mutants (light grey) are plotted alongside the wildtype enzyme (grey), and single mutant counterparts that were combined together for comparison. All single domain mutants shown are color coded from most negative (red) to most positive (blue). Data reported represents the average of four technical replicates, and error bars represent standard deviation from the mean. (A) Glucose equivalents released after AFEX cornstover hydrolysis for D3 CBM2a – D4 Cel5A combinatorial mutant. (B) Glucose equivalents released after AFEX cornstover hydrolysis for D4 CBM2a – D3 Cel5A combinatorial mutant. (C) Glucose equivalents released after AFEX cornstover hydrolysis for D2 CBM2a – D3 Cel5A combinatorial mutant. (D) Glucose equivalents released after crystalline cellulose − I hydrolysis for D3 CBM2a – D4 Cel5A combinatorial mutant. (E) Glucose equivalents released after crystalline cellulose − I hydrolysis for D4 CBM2a – D3 Cel5A combinatorial mutant. (F) Glucose equivalents released after crystalline cellulose – I hydrolysis for D2 CBM2a – D3 Cel5A combinatorial mutant.

When combining two oppositely supercharged domains as is the case in the third combinatorial construct D2 CBM2a – D3 Cel5A, activity is almost completely killed. On biomass, the D2 CBM2a – D3 Cel5A mutant displays low activity similar to its positively supercharged D3 CD. Even though the D2 CBM mutant was the most active individual construct on biomass, combination with the positively supercharged D3 CD led to upwards of 2-fold reductions in activity. This effect is even more drastic on crystalline cellulose, and on this substrate the combinatorial mutant shows little to no activity. There seems to be little synergism between the negatively charged CBM and positively charged CD. Both individual mutants have different ionization points, and different optimal pH where they are most active, translating to a combinatorial mutant with poor stability and activity. There is also the possibility for unfavorable intramolecular electrostatic interactions between oppositely charged domains that may perturb orientation of the CBM binding face and Cel5A active site with substrate. These effects were not observed in cell lysate assays likely due to the high concentration of salt and stabilizers like glycerol in the lysis buffer that would help keep the protein stable and screen unfavorable charged interactions. All purified enzyme assays reported utilized a minimal amount of salt in order to prevent charge screening that can mask interactions that result from supercharging.

## CONCLUSION

In this work we have successfully supercharged a family-5 endoglucanase Cel5A and its native family-2a carbohydrate binding module from the thermophilic microbe *Thermobifida fusca* in order to change surface charged interactions between enzyme and substrate. A total library size of 33 mutant constructs was created from computational CBM and CD designs with non-natural net charges. Characterization of soluble cell lysates for all 33 mutants and purified enzymes resulted in the following key conclusions: (i) hydrolytic activity is correlated with enzyme surface charge, (ii) supercharging only the CBM is more effective at improving catalytic activity, (iii) the optimal pH for biomass hydrolysis can be shifted through supercharging, and (iv) positive supercharging of the CBM can be used to improve thermostability in presence of cellulosic substrate. In addition to these conclusions, three key constructs were identified as being more active than the wildtype full length enzyme: (i) D2 CBM2a – WT Cel5A, (ii) D3 CBM2a – WT Cel5A, and (iii) D4 CBM2a – WT Cel5A. This is the first reported work in the field that has been able to successfully utilize supercharging approach to improve activity on both pretreated biomass and crystalline cellulose.

The three key CBM construct created in this work with better catalytic performance than the wildtype enzyme bare important outcomes with regards to sustainability and implementation into industrial biorefineries. The cost of using inefficient CAZymes in industrial biorefineries significantly limits the economic viability of biofuel production. The improved CBM constructs that show up to a 2-fold reduction in enzyme loading can result in up to a $0.57 reduction in cost per gallon of ethanol^20^ at normal processing temperatures. For the two positively supercharged CBM mutants with elevated optimal temperatures, even greater improvements to hydrolysis yield can translate to further economic cost reduction for final products. Lastly, modified pH optima for different CBM2a – Cel5A mutants has implications for fine-tuning biofuel processing conditions using either yeast that require acidic medium (pH 4.0 to 6.0)^55^ or bacteria like *E. coli* which require more neutral conditions^56^.

The effect of shifting enzyme surface charge closely resembles volcano plots used to find a Sabatier optimum when correlating activity to binding affinity^32–34^. In this scenario, results from previous studies^29,30^ as well as this current work suggest that enzyme binding to crystalline cellulose and cellulosic substrates is highly dependent on enzyme net charge. Utilizing GFP based pull-down binding assays we have shown that manipulating surface charge via protein supercharging can alter the binding affinity for these enzymes. Correlating these net charges to the activity for purified enzymes elucidates optimal net charges where activity peaks resembling a Sabatier optimum similar to those observed for cellulases in previous studies. The complex nature of substrate, as well as multifaceted interactions between solubilized enzyme and insoluble substrate limits net charge from being a one-dimensional predictor of binding affinity and improved activity. This can be observed when comparing optimal net charges between biomass and crystalline cellulose. There is clearly a benefit to positively supercharging enzymes to facilitate binding to a slightly negatively charged substrate like cellulose, but with a more complex substrate like lignocellulosic biomass that contains lignin and soluble xylan, the net charge optimum shifts and favors negative charges. Still with lignin present, positively charged CBM constructs showed improved activity on biomass at neutral pH’s likely due to the greater contribution to activity from the improved binding to cellulose by nature of their positive charges. This effect can be extended when analyzing activity at elevated temperatures, where both positively supercharged mutants exhibit elevated temperature optimums again due to their improved interactions with cellulose. Many of these trends do not hold for the peculiar D2 CBM2a – WT Cel5A construct. With a negatively supercharged CBM, this mutant exhibited improved biomass hydrolysis across a range of pH, as well as unexpectedly high activity on crystalline cellulose in a narrow pH range. This behavior seems to be the only outlier when comparing net charge to activity, and the cause for these improvements isn’t fully understood. It is likely that protein supercharging resulted in some structural anomalies that may alter the structure-function relationship of this mutant. These changes in overall structure may be impacting the overall solution stability, as well as normal CBM function such as how the planar aromatic residues align with glucopyranose rings within the dextran chain, substrate channeling into the Cel5A active site, adsorption to cellulose, desorption from cellulose, or penetration into the bulk substrate^26,57,58^. Elucidating the source for these improvements and understanding how supercharging may have altered the overall CBM architecture will require more detailed characterization in future work.

## Supporting information

SI Sequences Appendix

## ACKNOWLEDGEMENTS

This study was primarily supported by NSF CBET awards (1604421 and 1846797). ADC was supported by the Biotechnology Training Program fellowship that was funded by Rutgers University and the National Institute of General Medical Sciences of the National Institutes of Health under award number T32 GM135141. All genes used for expression of recombinant constructs were provided by the Department of Energy Joint Genome Institute (DOE-JGI) supported by the Community Science Program Gene Synthesis Award (CSP-503631 Syn). Finally, we would like to also thank Professor Rebecca Garlock-Ong (MTU) and Professor Bruce Dale (MSU) for kindly providing access to AFEX treated corn stover used in this study.

