## Supplementary material for "Supercharging carbohydrate-binding module alone enhances endocellulase thermostability, binding, and activity on cellulosic biomass": SI Appendix

**Number of SI pages in current PDF (SI-Supplementary Information)**: 13

**Number of SI figures in current PDF (SI-Supplementary Information): 5**

**Number of SI tables in current PDF (SI-Supplementary Information): 7**

**Number of SI files (including current PDF): 2**

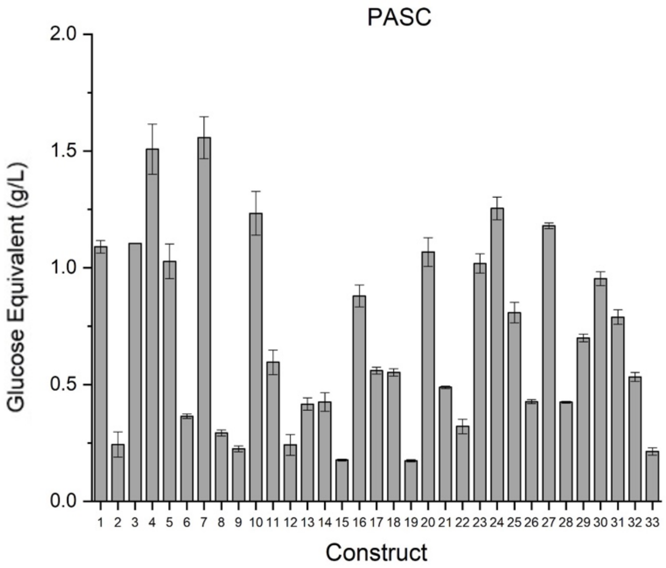


| Number | Construct Name | Number | Construct Name |
|---|---|---|---|
| 1 | Wild Type (WT) | 18 | D3 CBM2a – D2 Cel5A |
| 2 | WT CBM2a – D6 Cel5A | 19 | D3 CBM2a – D1 Cel5A |
| 3 | WT CBM2a – D4 Cel5A | 20 | D2 CBM2a – WT Cel5A |
| 4 | WT CBM2a – D3 Cel5A | 21 | D2 CBM2a – D6 Cel5A |
| 5 | WT CBM2a – D2 Cel5A | 22 | D2 CBM2a – D5 Cel5A |
| 6 | WT CBM2a – D1 Cel5A | 23 | D2 CBM2a – D4 Cel5A |
| 7 | D4 CBM2a – WT Cel5A | 24 | D2 CBM2a – D3 Cel5A |
| 8 | D4 CBM2a – D6 Cel5A | 25 | D2 CBM2a – D2 Cel5A |
| 9 | D4 CBM2a – D5 Cel5A | 26 | D2 CBM2a – D1 Cel5A |
| 10 | D4 CBM2a – D3 Cel5A | 27 | D1 CBM2a – WT Cel5A |
| 11 | D4 CBM2a – D2 Cel5A | 28 | D1 CBM2a – D6 Cel5A |
| 12 | D4 CBM2a – D1 Cel5A | 29 | D1 CBM2a – D5 Cel5A |
| 13 | D3 CBM2a – WT Cel5A | 30 | D1 CBM2a – D4 Cel5A |
| 14 | D3 CBM2a – D6 Cel5A | 31 | D1 CBM2a – D3 Cel5A |
| 15 | D3 CBM2a – D5 Cel5A | 32 | D1 CBM2a – D2 Cel5A |
| 16 | D3 CBM2a – D4 Cel5A | 33 | D1 CBM2a – D1 Cel5A |
| 17 | D3 CBM2a – D3 Cel5A | | |

**Figure S1. Activity of soluble cell lysates on phosphoric acid swollen cellulose (PASC).** All 33 constructs expressed as 200mL auto-induction cultures were pelleted and sonicated in buffer containing 20 mM sodium phosphate pH 7.4, 500 mL sodium chloride, and 20% (v/v) glycerol. 100 µL of cell lysate was incubated with 100 µL of PASC prepared as a 10 g/L slurry and incubated for 6 hours at 60 °C. Reducing sugar equivalents were estimated via DNS assay and compared to glucose standards. Data reported represents the average of four technical replicates and error bars represent standard deviation from the mean.

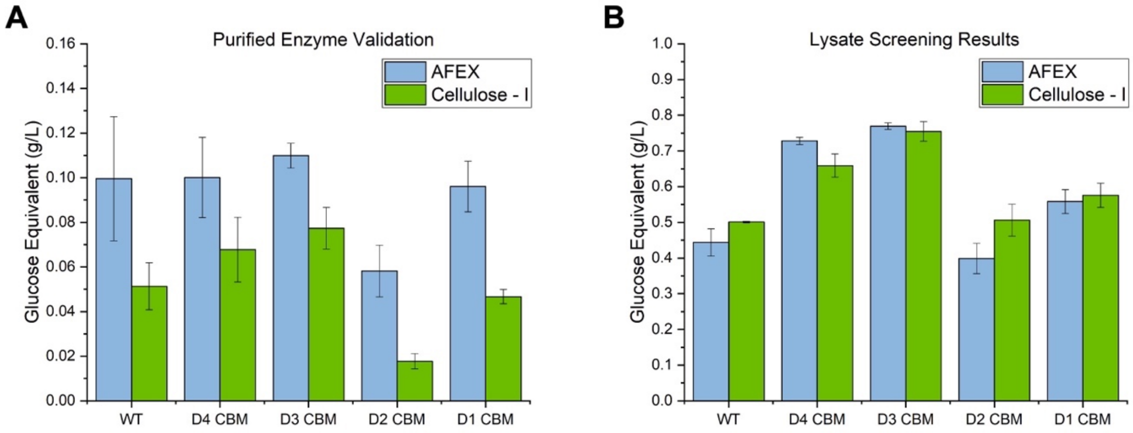


**Supplementary Figure 2. Purified enzyme assay to validate lysate screening results.** (A) Hydrolysis of AFEX corn stover (blue) and cellulose – I (green) by the wild type enzyme and all four CBM mutants (WT Cel5A CD) was conducted in the same cell lysis buffer (20 mM sodium phosphate, 500 mM NaCl, 20% glycerol, pH 7.4) that was used to lyse cells for lysate screening. A total of 2mg of substrate, either AFEX or cellulose – I was incubated with 120 nmol enzyme per gram of substrate in the presence of cell lysis buffer for 24 hours at 60 °C. Trends observed in (A) directly correlate to those observed for the same construct results reported in lysate screening depicted here in (B). The purified enzyme validation assay is in agreement with the results from lysate screening depicting both D3 CBM2a and D4 CBM2a as the best performing mutants, while D2 CBM2a is significantly hindered by the solution conditions, evident in both (A) and (B). Data reported represents the average of four technical replicates and error bars represent one standard deviation from the mean.

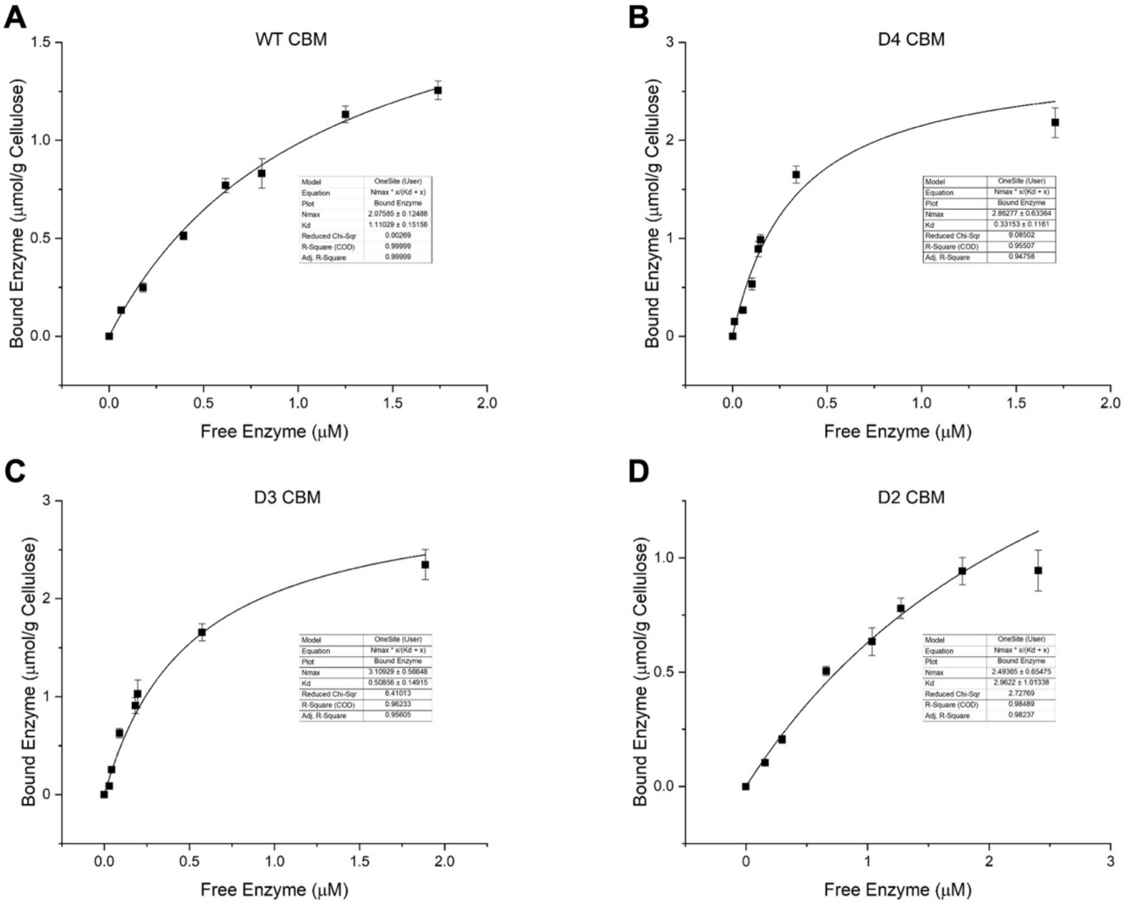


**Supplementary Figure 3. CBM-GFP full scale binding curves.** GFP was tagged to the (A) wild-type CBM2a, as well as three supercharged constructs, namely (B) D4 CBM2a, (C) D3 CBM2a, and (D) D2 CBM2a. Pull down binding assays were conducted with 1mg total Avicel PH-101 crystalline cellulose with protein concentrations ranging from 25 – 500 µg/mL in tandem with shaken/unshaken standards that contained no cellulose. All assays were conducted in 0.2mL round bottom microplates (Greiner Bio-One) , incubated at room temperature (25°C) for one hour with 5 RPM end over end mixing (unshaken standards incubated on lab bench), and 100 µL of the binding supernatant was aliquoted into opaque flat bottom microplates for measuring residual fluorescence after binding with cellulose. Results were plotted with Origin software, and data was fit to a one-site Langmuir model resulting in the trendline and fit parameters displayed. All data reported is an average of six technical replicates, and error bars represent standard deviation from the mean.

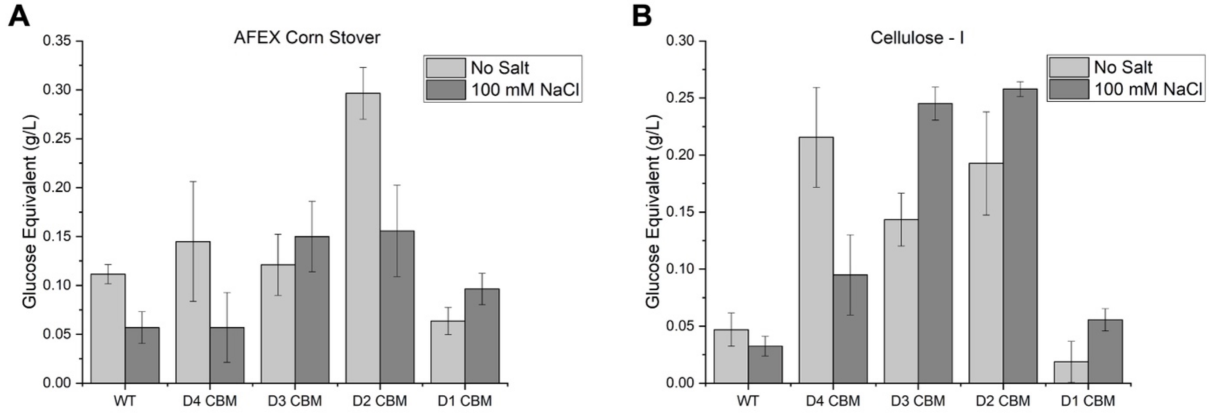


**Supplementary Figure 4. The addition of salt to screen charged interactions leads to substrate dependent alteration in supercharged enzyme activity.** A total of 4 mg of (A) AFEX corn stover or (B) cellulose – I was hydrolyzed by 120 nmol of wildtype, or CBM mutant enzyme per gram of substrate with (dark grey) and without (light grey) 100 mM NaCl. Reaction mixtures were incubated for 24 hours at 60 °C and reducing sugar equivalents measured by DNS reducing sugar assay. Data reported represents the average of four technical replicates and error bars represent on standard deviation from the mean. The addition of salt has a pronounced impact on catalytic activity depending on the substrate tested, with up to a possible 5-fold increase in activity observed compared to wildtype for the D2 CBM mutant when salt is added.

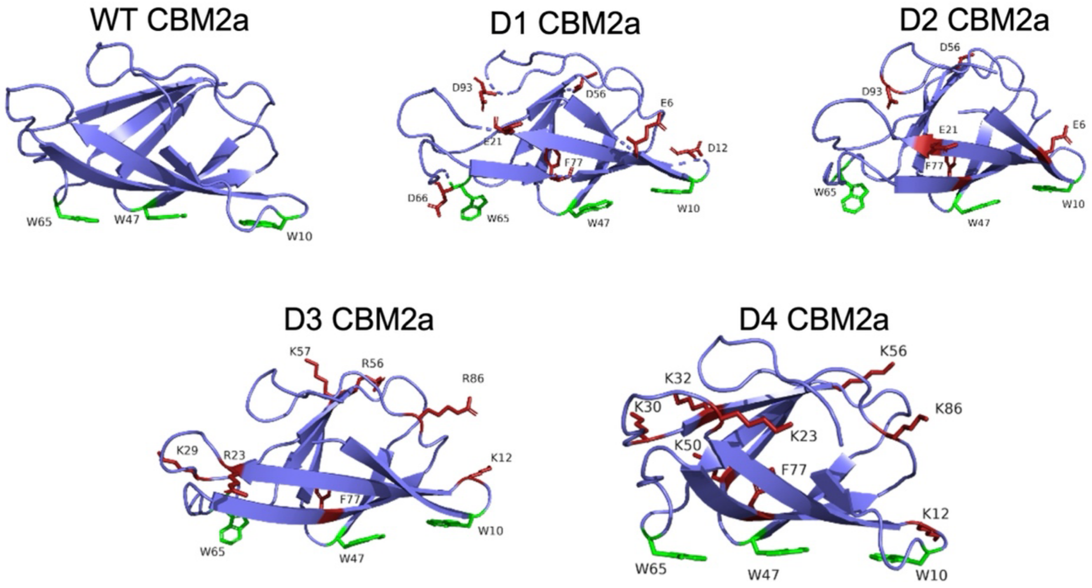


**Supplementary Figure 5. CBM2a binding module homology models for wildtype enzyme (WT) and four supercharged designs (D1 – D4)**. No solved crystal structure has been published for CBM2a from *T. fusca*, so a homology model for WT CBM2a was constructed using Rosetta CM and served as the basis for the supercharged domain homology models. Side chains of key amino acid resides on the CBM surface have been highlighted and labelled. Planar aromatic residues (W10, W47, W65) that modulate CBM binding to cellulose are highlighted in green. All mutated residues are highlighted in red and labelled for each design. Protein structure images were produced and analyzed in PyMOL.

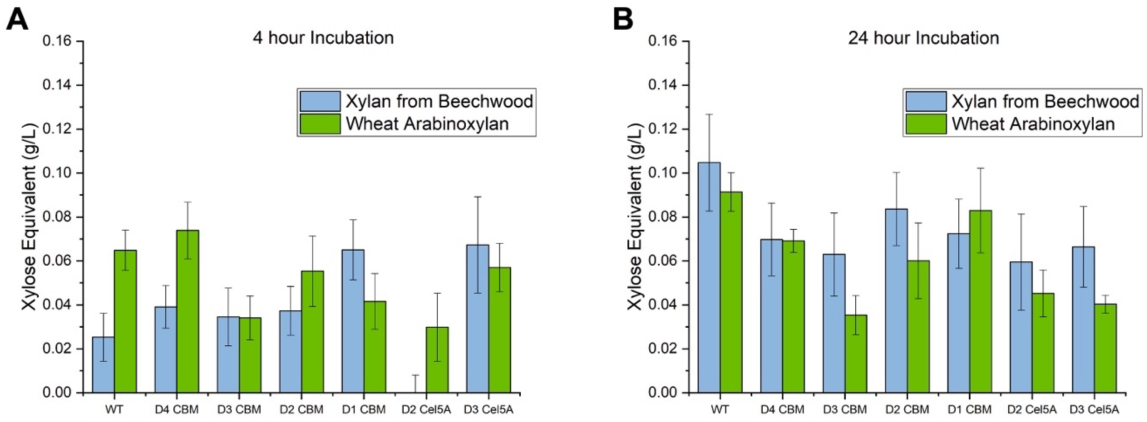


**Supplementary Figure 6. Hydrolysis of soluble Xylan substrates depicts little improvement in catalytic activity on xylan compared to wildtype.** Enzyme activity assays for the wild type enzyme, four CBM mutant constructs (WT Cel5A), and two mutant Cel5A constructs (D2/D3 Cel5A, WT CBM2a) were conducted to deconvolute the observed enzyme activity on pretreated biomass. Xylan from beechwood (blue) and wheat arabinoxylan (green) were prepared by dissolving each substrate in boiling deionized water as 10 g/L stock solutions. Hydrolysis assays were conducted by incubating 1 mg of substrate with 60 nmol of enzyme per gram of substrate at 65 °C for (A) 4 hours and (B) 24 hours. Reducing sugar concentration was estimated via DNS reducing sugar assay and compared to xylose standards. Data reported represents the average of four technical replicates and error bars represent standard deviation from the mean.

| CBM Design | Mutation |
|---|---|
| D1 | K6E, N12D, R21E, N56D, N66D, N93D |
| D2 | K6E, R21E, N56D, N93D |
| D3 | N12K, D23R,S29K, N56R, S57K, E86R |
| D4 | N12K, D23K, Q30K, E32K, Q50K, N56K, E86K |

**Supplementary Table T1. Mutations on wildtype CBM2a to generate supercharged mutants.** Surface residues identified using Rosetta macromolecular software were mutated to either a negatively charged amino acid (D, E) or positively charged amino acid (R, K) to supercharge the domain and obtain a desired net charge. This table lists the mutations necessary to generate the indicated mutant from wildtype CBM2a.

| CD Design | Mutations |
|---|---|
| D1 | R128E, K131E, N145D, K177E, N195D, R197E, Q205E, Q245E, R246E, Q282E, N313D, N316D, R332E, N334D, N337D, R340E, Q367E, R371E, K380E, Q419E, Q426E |
| D2 | R128E, K131E, N145D, K177E, N195D, R197E, Q205E, Q245E, R246E, Q282E, N313D, N316D, N337D, R340E, Q367E, R371E, K380E, Q419E, Q426E |
| D3 | H143R, Q245R, S268R, Q282R, E341K, E344K, Q367K, D374K, E378R, Q419K |
| D4 | H143R, Q245R, S268R, Q282R,N334K, E341K, E344K, Q367K, D374K, E378R, F392K, Q419K |
| D5 | E127K, E142K, N145K, D166K, D170K, D190K, N195K, D201K, Q205K, D208K, D234K, Q245K, E275K, Q282K, D286K, N313K, N316K, D333K, N337K, E341K, E344K, Q367K, D370K, D374K, E378K, Q419K, Q426K |
| D6 | E127K, E142K, N145K, D166K, D170K, D190K, N195K, D201K, Q205K, D208K, D234K, Q245K, E275K, Q282K, D286K, N313K, N316K, D333K, N334K, N337K, E341K, E344K, D361K, Q367K, D370K, D374K, E378K, Q419K, Q426K |

**Supplementary Table T2. Mutations on wildtype Cel5A to generate supercharged mutants.** Surface residues identified using Rosetta macromolecular software were mutated to either a negatively charged amino acid (D, E) or positively charged amino acid (R, K) to supercharge the domain and obtain a desired net charge. This table lists the mutations necessary to generate the indicated mutant from wildtype Cel5A.

| Design No. | Construct Name | Net Charge | pNPC (%) | Standard Deviation | AFEX corn stover (g/L) | Standard Deviation | Cellulose-I (g/L) | Standard Deviation |
|---|---|---|---|---|---|---|---|---|
| 1 | WT CBM2a - WT Cel5A | -8 | 4.430 | 0.4495 | 0.444 | 0.0382 | 0.501 | 0.0022 |
| 2 | WT CBM2a - D6 Cel5A | 38 | 0.047 | 0.0963 | 0.549 | 0.1199 | 0.229 | 0.0313 |
| 3 | WT CBM2a - D4 Cel5A | 7 | 0.389 | 0.0914 | 0.647 | 0.0243 | 0.463 | 0.0002 |
| 4 | WT CBM2a - D3 Cel5A | 5 | 1.404 | 0.2013 | 0.813 | 0.1124 | 0.690 | 0.1163 |
| 5 | WT CBM2a - D2 Cel5A | -35 | 0.550 | 0.0171 | 0.342 | 0.0418 | 0.496 | 0.0239 |
| 6 | WT CBM2a - D1 Cel5A | -38 | 0.007 | 0.0174 | 0.287 | 0.0189 | 0.226 | 0.0147 |
| 7 | D4 CBM2a - WT Cel5A | 4 | 2.770 | 0.3062 | 0.728 | 0.0099 | 0.746 | 0.0325 |
| 8 | D4 CBM2a - D6 Cel5A | 50 | 0.024 | 0.0182 | 0.328 | 0.0076 | 0.240 | 0.0299 |
| 9 | D4 CBM2a - D5 Cel5A | 47 | 0.024 | 0.0176 | 0.318 | 0.0159 | 0.211 | 0.0205 |
| 10 | D4 CBM2a - D3 Cel5A | 17 | 0.568 | 0.2009 | 0.585 | 0.0220 | 0.659 | 0.0236 |
| 11 | D4 CBM2a - D2 Cel5A | -23 | 0.055 | 0.0300 | 0.404 | 0.0506 | 0.411 | 0.0119 |
| 12 | D4 CBM2a - D1 Cel5A | -26 | 0.193 | 0.0542 | 0.327 | 0.0196 | 0.233 | 0.0149 |
| 13 | D3 CBM2a - WT Cel5A | 2 | 28.167 | 0.2129 | 0.769 | 0.0094 | 0.755 | 0.0274 |
| 14 | D3 CBM2a - D6 Cel5A | 48 | 0.391 | 0.0490 | 0.548 | 0.0345 | 0.394 | 0.0296 |
| 15 | D3 CBM2a - D5 Cel5A | 45 | 0.144 | 0.0967 | 0.263 | 0.0299 | 0.202 | 0.0095 |
| 16 | D3 CBM2a - D4 Cel5A | 17 | 0.183 | 0.0959 | 0.490 | 0.0372 | 0.552 | 0.0253 |
| 17 | D3 CBM2a - D3 Cel5A | 15 | 0.153 | 0.0683 | 0.280 | 0.0320 | 0.269 | 0.0099 |
| 18 | D3 CBM2a - D2 Cel5A | -25 | 0.147 | 0.0581 | 0.404 | 0.0438 | 0.366 | 0.0116 |
| 19 | D3 CBM2a - D1 Cel5A | -28 | 0.135 | 0.0492 | 0.199 | 0.0116 | 0.155 | 0.0169 |
| 20 | D2 CBM2a - WT Cel5A | -12 | 7.163 | 0.5031 | 0.399 | 0.0424 | 0.506 | 0.0446 |
| 21 | D2 CBM2a - D6 Cel5A | 34 | 0.158 | 0.0326 | 0.397 | 0.0588 | 0.375 | 0.0335 |
| 22 | D2 CBM2a - D5 Cel5A | 31 | 0.167 | 0.0392 | 0.378 | 0.0388 | 0.304 | 0.0334 |
| 23 | D2 CBM2a - D4 Cel5A | 3 | 0.692 | 0.0625 | 0.340 | 0.0280 | 0.420 | 0.0107 |
| 24 | D2 CBM2a - D3 Cel5A | 1 | 2.620 | 0.1796 | 0.492 | 0.0366 | 0.613 | 0.0143 |
| 25 | D2 CBM2a - D2 Cel5A | -39 | 0.092 | 0.1261 | 0.373 | 0.0234 | 0.406 | 0.0279 |
| 26 | D2 CBM2a - D1 Cel5A | -42 | -0.079 | 0.1193 | 0.308 | 0.0278 | 0.301 | 0.0252 |
| 27 | D1 CBM2a - WT Cel5A | -14 | 4.346 | 1.3464 | 0.558 | 0.0333 | 0.575 | 0.0339 |
| 28 | D1 CBM2a - D6 Cel5A | 32 | 0.050 | 0.1277 | 0.470 | 0.0184 | 0.406 | 0.0241 |
| 29 | D1 CBM2a - D5 Cel5A | 29 | 0.485 | 0.7507 | 0.734 | 0.0439 | 0.668 | 0.0469 |
| 30 | D1 CBM2a - D4 Cel5A | 1 | 0.174 | 0.1642 | 0.630 | 0.0125 | 0.589 | 0.0589 |
| 31 | D1 CBM2a - D3 Cel5A | -1 | 0.258 | 0.1795 | 0.451 | 0.0149 | 0.368 | 0.0426 |
| 32 | D1 CBM2a - D2 Cel5A | -41 | -0.019 | 0.1635 | 0.337 | 0.0304 | 0.285 | 0.0349 |
| 33 | D1 CBM2a - D1 Cel5A | -44 | 0.032 | 0.1250 | 0.261 | 0.0122 | 0.183 | 0.0087 |
| N/A | T7 Shuffle Negative Control | N/A | 0 | | 0 | | 0 | |

**Supplementary Table T3. Tabulated summary of lysate screening results.** Summary of all 33 constructs, net charges, and absolute activities measured on pNPC, AFEX corn stover and cellulose – I. Enzymes highlighted in green reported higher activity than wildtype enzyme on at least two substrates tested. This data is plotted in **Fig. 2** of the main manuscript. Net charges were estimated by counting the number of charged amino acid residues in each enzyme's sequence.

| | | AFEX CS | | | | | |
|---|---|---|---|---|---|---|---|
| **Construct 1** | **Construct 2** | **pH 4.5** | **pH 5.0** | **pH 5.5** | **pH 6.0** | **pH 6.5** | **pH 7.0** |
| **WT_WT** | D4_WT | 0.0055 | 0.1657 | 0.4529 | 0.0037 | 0.0034 | 0.0056 |
| | D3_WT | 0.0071 | 0.0002 | 0.2612 | 0.1150 | 0.0007 | 0.0001 |
| | D2_WT | 0.0002 | 0.0003 | 0.0076 | 0.1183 | 0.0010 | 0.0103 |
| | D1_WT | 0.0055 | 0.0146 | 0.2464 | 0.0029 | 0.0156 | 0.0541 |
| | WT_D2 | 0.1147 | 0.0128 | 0.0005 | 0.0001 | 0.0147 | 0.0027 |
| | WT_D3 | 0.0091 | 0.0340 | 0.0383 | 0.0148 | 0.3323 | 0.3057 |
| **D4_WT** | D3_WT | 0.0102 | 0.0077 | 0.1500 | 0.0078 | 0.0035 | 0.0001 |
| | D2_WT | 0.0006 | 0.0026 | 0.0079 | 0.0232 | 0.0343 | 0.0315 |
| | D1_WT | 0.0298 | 0.1147 | 0.1386 | 0.1570 | 0.0079 | 0.0082 |
| | WT_D2 | 0.1288 | 0.0559 | 0.0002 | 0.0001 | 0.0001 | 0.0005 |
| | WT_D3 | 0.0631 | 0.0251 | 0.0199 | 0.0005 | 0.0009 | 0.0402 |
| **D3_WT** | D2_WT | 0.0042 | 0.0006 | 0.0108 | 0.2275 | 0.0023 | 0.0045 |
| | D1_WT | 0.4047 | 0.0296 | 0.1331 | 0.0084 | 0.0006 | 0.0007 |
| | WT_D2 | 0.0149 | 0.0001 | 0.0006 | 0.0001 | 0.0000 | 0.0001 |
| | WT_D3 | 0.3945 | 0.0190 | 0.0047 | 0.0033 | 0.0001 | 0.0020 |
| **D2_WT** | D1_WT | 0.0009 | 0.0034 | 0.0063 | 0.0411 | 0.4327 | 0.0044 |
| | WT_D2 | 0.0002 | 0.0001 | 0.0020 | 0.0007 | 0.0007 | 0.0000 |
| | WT_D3 | 0.0001 | 0.0032 | 0.0255 | 0.0006 | 0.0160 | 0.0031 |
| **D1_WT** | WT_D2 | 0.0246 | 0.0038 | 0.0005 | 0.0000 | 0.0007 | 0.0104 |
| | WT_D3 | 0.4737 | 0.3010 | 0.0247 | 0.0009 | 0.0001 | 0.1567 |
| **WT_D2** | WT_D3 | 0.0283 | 0.0138 | 0.0168 | 0.0143 | 0.0043 | 0.0054 |

**Supplementary Table T4. Student's T-test p-values for the comparison of purified enzyme activity on AFEX cornstover at different pH.** The raw data corresponding to these statistics is reported in **Fig. 3A** and **Fig. 3B** in the main manuscript.

| | | Cellulose I | | | | | |
|---|---|---|---|---|---|---|---|
| Construct 1 | Construct 2 | pH 4.5 | pH 5.0 | pH 5.5 | pH 6.0 | pH 6.5 | pH 7.0 |
| WT_WT | D4_WT | 0.0030 | 0.0092 | 0.0042 | 0.0028 | 0.0002 | 0.0122 |
| | D3_WT | 0.0012 | 0.0007 | 0.0069 | 0.0001 | 0.0001 | 0.0001 |
| | D2_WT | 0.0008 | 0.0002 | 0.0036 | 0.2598 | 0.3505 | 0.4871 |
| | D1_WT | 0.0193 | 0.0067 | 0.2068 | 0.0022 | 0.0413 | 0.0090 |
| | WT_D2 | 0.0113 | 0.0439 | 0.0000 | 0.0001 | 0.0109 | 0.0008 |
| | WT_D3 | 0.0040 | 0.0277 | 0.0150 | 0.0191 | 0.0061 | 0.0081 |
| D4_WT | D3_WT | 0.2491 | 0.0085 | 0.0178 | 0.3966 | 0.0587 | 0.0013 |
| | D2_WT | 0.0019 | 0.0059 | 0.0245 | 0.0049 | 0.0009 | 0.0178 |
| | D1_WT | 0.0463 | 0.2449 | 0.0136 | 0.0003 | 0.0005 | 0.0061 |
| | WT_D2 | 0.0112 | 0.0113 | 0.0005 | 0.0003 | 0.0005 | 0.0009 |
| | WT_D3 | 0.0213 | 0.4324 | 0.0424 | 0.2184 | 0.0019 | 0.1049 |
| D3_WT | D2_WT | 0.0022 | 0.1195 | 0.1505 | 0.0029 | 0.0000 | 0.0001 |
| | D1_WT | 0.0492 | 0.0012 | 0.0080 | 0.0004 | 0.0000 | 0.0001 |
| | WT_D2 | 0.0048 | 0.0001 | 0.0017 | 0.0000 | 0.0000 | 0.0000 |
| | WT_D3 | 0.0931 | 0.0120 | 0.0129 | 0.1313 | 0.0303 | 0.0000 |
| D2_WT | D1_WT | 0.1314 | 0.0043 | 0.0056 | 0.0372 | 0.0000 | 0.0002 |
| | WT_D2 | 0.0011 | 0.0000 | 0.0011 | 0.0024 | 0.0001 | 0.0029 |
| | WT_D3 | 0.0087 | 0.0136 | 0.1239 | 0.0142 | 0.0188 | 0.0009 |
| D1_WT | WT_D2 | 0.0244 | 0.0009 | 0.0114 | 0.0018 | 0.0006 | 0.0184 |
| | WT_D3 | 0.1115 | 0.2120 | 0.0033 | 0.0099 | 0.0102 | 0.0014 |
| WT_D2 | WT_D3 | 0.0074 | 0.0260 | 0.0013 | 0.0023 | 0.0086 | 0.0010 |

**Supplementary Table T5. Student's T-test p-values for the comparison of purified enzyme activity on cellulose – I at different pH.** The raw data corresponding to these statistics is reported in **Fig. 3C** and **Fig. 3D** in the main manuscript.

| Construct 1 | Construct 2 | 55 C | 60 C | 65C | 70C | 75C | 80C |
|---|---|---|---|---|---|---|---|
| WT_WT | D4_WT | 0.0085 | 0.0076 | 0.0002 | 0.0000 | 0.0004 | 0.0018 |
| | D3_WT | 0.0124 | 0.0004 | 0.0001 | 0.0000 | 0.0001 | 0.0005 |
| | D2_WT | 0.1164 | 0.0899 | 0.3876 | 0.0011 | 0.0008 | 0.2434 |
| | D1_WT | 0.0034 | 0.0009 | 0.0003 | 0.0000 | 0.0015 | 0.0000 |
| | WT_D2 | 0.0017 | 0.0004 | 0.0000 | 0.0000 | 0.0001 | 0.0002 |
| | WT_D3 | 0.4618 | 0.0206 | 0.1034 | 0.0001 | 0.0008 | 0.2588 |
| D4_WT | D3_WT | 0.1589 | 0.1589 | 0.0001 | 0.0006 | 0.0006 | 0.0365 |
| | D2_WT | 0.0002 | 0.0002 | 0.0005 | 0.0000 | 0.0011 | 0.0007 |
| | D1_WT | 0.4567 | 0.4567 | 0.0059 | 0.1614 | 0.0272 | 0.0536 |
| | WT_D2 | 0.0113 | 0.0113 | 0.0000 | 0.0000 | 0.0000 | 0.0000 |
| | WT_D3 | 0.0175 | 0.0175 | 0.0027 | 0.0010 | 0.0008 | 0.0100 |
| D3_WT | D2_WT | 0.0002 | 0.0000 | 0.0002 | 0.0001 | 0.0002 | 0.0093 |
| | D1_WT | 0.0523 | 0.0007 | 0.0009 | 0.0005 | 0.0016 | 0.0019 |
| | WT_D2 | 0.0002 | 0.0005 | 0.0000 | 0.0001 | 0.0002 | 0.0192 |
| | WT_D3 | 0.0034 | 0.0001 | 0.0000 | 0.0001 | 0.0000 | 0.0024 |
| D2_WT | D1_WT | 0.0012 | 0.0002 | 0.0003 | 0.0003 | 0.0060 | 0.0020 |
| | WT_D2 | 0.0252 | 0.0000 | 0.0000 | 0.0000 | 0.0001 | 0.0001 |
| | WT_D3 | 0.1998 | 0.3992 | 0.1497 | 0.0040 | 0.0032 | 0.0070 |
| D1_WT | WT_D2 | 0.0011 | 0.0006 | 0.0000 | 0.0000 | 0.0003 | 0.0009 |
| | WT_D3 | 0.0093 | 0.0003 | 0.0001 | 0.0000 | 0.0478 | 0.0007 |
| WT_D2 | WT_D3 | 0.0123 | 0.0001 | 0.0000 | 0.0000 | 0.0000 | 0.0000 |

**Supplementary Table T6. Student's T-test p-values for the comparison of purified enzyme activity on pNP cellobiose incubated at different temperatures.** The raw data corresponding to these statistics is reported in **Fig. 5A** in the main manuscript.

| Construct 1 | Construct 2 | 55 C | 60 C | 65C | 70C | 75C | 80C |
|---|---|---|---|---|---|---|---|
| WT_WT | D4_WT | 0.0085 | 0.2810 | 0.0069 | 0.0088 | 0.0004 | 0.0433 |
| | D3_WT | 0.0124 | 0.3996 | 0.0032 | 0.0025 | 0.0119 | 0.0412 |
| | D2_WT | 0.1164 | 0.0624 | 0.0742 | 0.0188 | 0.1196 | 0.0084 |
| | D1_WT | 0.0034 | 0.0093 | 0.0101 | 0.1352 | 0.0052 | 0.0021 |
| | WT_D2 | 0.0017 | 0.0029 | 0.0055 | 0.0337 | 0.0038 | 0.0000 |
| | WT_D3 | 0.4618 | 0.3068 | 0.0204 | 0.0043 | 0.4668 | 0.2588 |
| D4_WT | D3_WT | 0.0222 | 0.0222 | 0.3532 | 0.0002 | 0.1073 | 0.2762 |
| | D2_WT | 0.3050 | 0.3050 | 0.0009 | 0.0030 | 0.0000 | 0.0138 |
| | D1_WT | 0.0003 | 0.0003 | 0.0133 | 0.0000 | 0.0000 | 0.0106 |
| | WT_D2 | 0.0000 | 0.0000 | 0.0132 | 0.0000 | 0.0000 | 0.0030 |
| | WT_D3 | 0.0031 | 0.0031 | 0.0027 | 0.1566 | 0.0001 | 0.1038 |
| D3_WT | D2_WT | 0.3760 | 0.1762 | 0.0054 | 0.0004 | 0.0123 | 0.0409 |
| | D1_WT | 0.0004 | 0.0186 | 0.0002 | 0.0000 | 0.0049 | 0.0076 |
| | WT_D2 | 0.0000 | 0.0027 | 0.0003 | 0.0000 | 0.0020 | 0.0036 |
| | WT_D3 | 0.0047 | 0.3399 | 0.0013 | 0.0014 | 0.0106 | 0.1000 |
| D2_WT | D1_WT | 0.0081 | 0.0003 | 0.0022 | 0.0002 | 0.0002 | 0.0292 |
| | WT_D2 | 0.0208 | 0.0001 | 0.0006 | 0.0000 | 0.0005 | 0.0055 |
| | WT_D3 | 0.1055 | 0.1480 | 0.2192 | 0.0008 | 0.0041 | 0.4203 |
| D1_WT | WT_D2 | 0.4008 | 0.0003 | 0.0088 | 0.0008 | 0.0568 | 0.0044 |
| | WT_D3 | 0.0007 | 0.0074 | 0.0001 | 0.0001 | 0.0006 | 0.0757 |
| WT_D2 | WT_D3 | 0.0014 | 0.0049 | 0.0001 | 0.0000 | 0.0023 | 0.0282 |

**Supplementary Table T7. Student's T-test p-values for the comparison of purified enzyme activity on cellulose-I incubated at different temperatures.** The raw data corresponding to these statistics is reported in **Fig. 5B** in the main manuscript.
